# Divergence between transcriptomes and chromatin accessibility during differentiation from a bipotential progenitor cell population to erythroblasts and megakaryocytes

**DOI:** 10.1101/2025.06.30.662383

**Authors:** Tejaswini Mishra, Belinda M. Giardine, Christapher S. Morrissey, Cheryl A. Keller, Elisabeth F. Heuston, Stacie M. Anderson, Vikram R. Paralkar, Maxim Pimkin, Mitchell J. Weiss, David M. Bodine, Ross C. Hardison

## Abstract

Changes in gene expression drive differentiation along distinct cell lineages, and these shifts in gene expression are associated with alterations in chromatin accessibility and modifications reflecting activation or repression. We used deep sequencing of polyA+ RNA to map the transcriptomes of the megakaryocyte-erythroid progenitor (MEP) and cells of its two daughter lineages, erythroblasts (ERY) and megakaryocytes (MEG) in mice to reveal insights into differentiation. Transcriptome comparisons revealed that MEPs already expressed much of the MEG program while continuing to express genes associated with parallel myeloid lineages. By contrast, ERY underwent an extensive program of gene induction along with repression of pan-hematopoietic and MEG genes. Maps of transcription factor (TF) occupancy also indicated distinct modes of regulation for the MEG and ERY programs, with MEG genes preferentially occupied by hematopoietic TFs in multipotent progenitors and continued occupancy post-commitment, in contrast to erythroid genes that were primarily occupied in committed ERY. Previous work had indicated a surprising discordance in the clustering of MEP with other hematopoietic cell types by RNA-seq *versus* chromatin states. We combined the differential expression data with chromatin accessibility across blood cell types to identify trends that contribute to this discordance. Specifically, candidate cis-regulatory elements (cCREs) in some ERY-specific genes were precociously actuated in the bipotential cell populations, and some other genes were expressed in both the MEP population and MEG but their cCREs have less chromatin accessibility in MEP. This discordance in cell type clustering by different modalities of functional genomics may reflect the different contributions of subpopulations in the MEP to the different modalities measured.

## Introduction

Cell fate decisions are executed via lineage-specific gene transcription (1), but our understanding of how these processes are initiated and maintained is incomplete. Differentiation of hematopoietic stem cells into multipotential progenitors and lineage-specified cells is an excellent system for investigating the transcriptional and epigenomic landscapes during commitment and maturation (2, 3) because stem and progenitor cell populations (4, 5) can be purified using cell surface markers and the cell lineages produced by each progenitor cell type are well-characterized (6). A particularly interesting cell fate decision is the differentiation of megakaryocytes (MEG) and erythroblasts (ERY) from a population of megakaryocyte-erythroid progenitor cells (MEP). This bifurcating differentiation stage is remarkable for the striking differences in the progeny cells. Committed proerythroblasts amplify by cell division (7, 8) and mature into erythrocytes with a remodeled cytoskeleton that contain large amounts of hemoglobin to facilitate oxygen and carbon dioxide transport (9). In contrast, committed megakaryoblasts undergo many rounds of DNA replication without cell division to generate polyploid MEG (10, 11) that are the source for platelet biogenesis (12). Despite the differences in maturation and function of ERY and MEG, these cell lineages can be derived from a common bipotential progenitor population, the MEP (13, 14).

The differentiation of MEG and ERY has been studied by multiple transcriptomic (15, 16) and functional genomic (17–24) approaches in mouse and human. These studies revealed that differentiation of both cell types depends on shared hematopoietic TFs including GATA1, GATA2, and TAL1 (25, 26), but these TFs exhibit distinctly different patterns of occupancy in the two lineages (17, 19, 20, 27), and the chromatin state maps in differentially expressed loci differ between the two lineages (21, 23, 24). In addition, ERY and MEG produce distinct TFs that facilitate their differentiation (18). For example, ETS factors such ERG and FLI1 are expressed preferentially in MEG and may be involved in establishing MEG-specific patterns of occupancy by GATA factors and TAL1. KLF1, an ERY-specific TF, is associated with some GATA1- and TAL1-bound DNA segments in erythroblasts (19, 28, 29). Furthermore, epigenomics studies using total RNA preparations and chromatin accessibility have indicated divergent modes of regulation in the two lineages (22). These studies of the population of hematopoietic stem cells termed LSK (lin-, Sca1+, Kit+) and the common myeloid progenitor (CMP) differentiating to MEG and ERY showed that MEG retained a transcriptome similar to that of the progenitor cell types whereas the transcriptomes of ERY reflected extensive lineage-specific induction and repression.

An open question about this bifurcation in differentiation is the relationship of MEP to other blood cell types. The placement of the population of bipotential MEP with other cell types differs depending on whether the distance measures are based on features of chromatin structure or the levels of stable RNA (Figure 1A). This discordance in the groupings has been reported in multiple studies of chromatin accessibility and histone modification along with RNA across many mouse blood cell types, including both purified hematopoietic stem and progenitor cell populations and mature cells from each lineage (22, 23, 30) (Supporting Information Figure S1). Specifically, MEP groups closely to erythroid cells when comparing patterns of chromatin accessibility determined by the assay for transposase accessible chromatin followed by sequencing (ATAC-seq) (22, 23) and in comparisons of profiles of the histone modification H3K4me1 (30). In contrast, MEP groups more closely to megakaryocytic cells and multilineage progenitors when examining stable RNA. A similar discordance in the groupings of cell types including MEP also was observed for human blood cells (31). These differences in groupings suggest some dissociation between chromatin profiles and transcriptome profiles around the stage of the bifurcation of MEP to MEG and ERY, but a more specific explanation has not yet been established.

**Figure 1.**
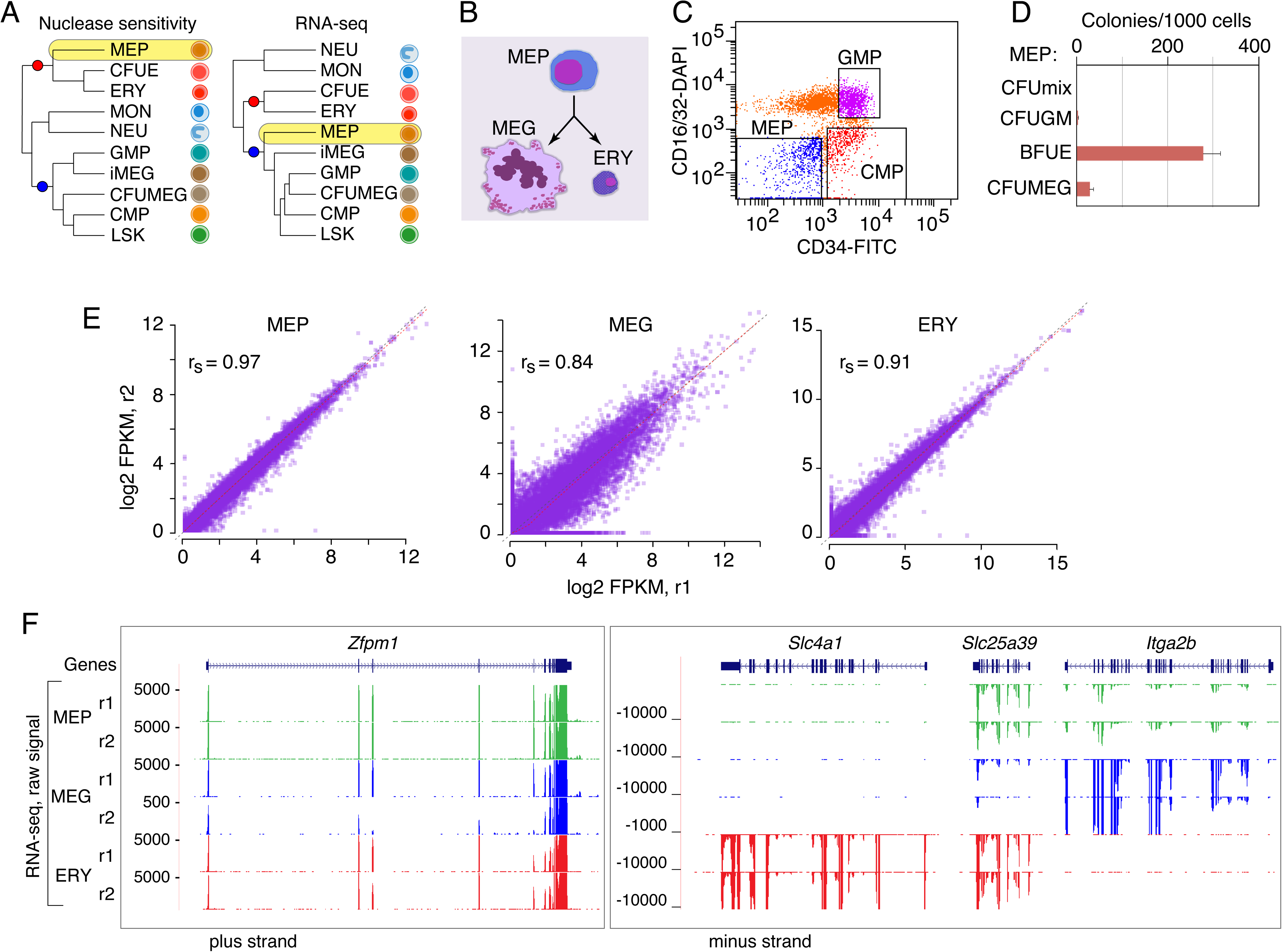
Discordant groupings and RNA-sequencing on MEP, MEG and ERY. (A) Consensus trees of relatedness among blood cell types based on chromatin accessibility (left) or transcriptome (right) distances, summarizing published results (22, 23, 30). The placement of MEP is highlighted in yellow, and nodes that differ in the trees are marked with red and blue circles. (B) Relationship among cell types for which polyA+ RNA transcriptome profiling was performed. (C) Results of FACS to purify the megakaryocyte-erythroid progenitor population using CD34 and CD16/32 markers. MEPs are CD34^low^ and CD16/32(−). (D) Barplot showing the frequency of formation of colonies of distinct cell types from the purified MEP population. (E) Scatter plots of the normalized log2 FPKMs measured for RNA from the 22,977 RefSeq genes, comparing the two replicates for MEP, MEG and ERY, along with the Spearman’s correlation coefficient. (E) Signal profiles for the raw polyA+ RNA-seq signal tracks from ERY, MEG and MEP, showing only the data on the transcribed strand (synonymous with the RNA). The diagram for the *Zfpm1* gene (left), which is expressed in all three cell types, covers a 64kb interval, chr8:124,802,001-124,866,000 on the mm9 genome assembly. The three closely linked, differentially expressed genes *Slc4a1*, *Slc25a39*, and *Itga2b*, are shown using Multi-Region view (right) to focus on the intervals encompassing each, specifically chr11:102207001-102230000, 102264001-102270000, and 102313001-102333000, respectively.

Transcriptome profiling in single cells across hematopoietic differentiation in mouse and human revealed that the multi-potent and bi-potent progenitor cells are not homogeneous cell populations, but rather they represent major transitional stages along a largely continuous trajectory of differentiation (32–34). The human MEP population has been shown to consist of at least three sub-populations, all of which have a component of bipotential cells. A majority of one subpopulation is bipotential cells, and the other two subpopulations are biased toward either erythroid or megakaryocytic differentiation (34, 35). Identification of bipotential MEP is challenging for many reasons, including the much higher proliferative potential of ERY than MEG and the fact that MEG and ERY require different culture conditions, which confounds assays for both cell types simultaneously (36). The single cell data suggest a common ancestor for progenitor cells committed to the erythroid and megakaryocytic lineages, but since there is no culture system that can produce mixed erythroid and megakaryocytic colonies, the common ancestor should be considered a model rather than a demonstrated cell type. Indeed, controversy persists over which purification schemes are best for MEP in mouse and human (36), but some methods for isolating MEP have been widely used (37).

We reasoned that some of the controversy about the MEP bifurcation in blood cell differentiation could arise from lower coverage RNA-sequencing in the rare progenitor cell populations and especially in the sparse coverage for single cell assays. Such low coverage could lead to missing some informative transcripts. To overcome the limitations of sparse coverage, we turned to deeply sequenced, polyA+ transcriptomes from mouse MEP, ERY, and MEG that had been previously analyzed for long noncoding transcripts in these lineages (38). The experiments generating these data sets analyzed transcriptomes of MEP, MEG, and ERY using directional RNA-sequencing (RNA-seq) to a very deep coverage, over 100 million reads per replicate (38). Thus, these data should reveal even rare transcripts. In this paper, we identify the transcriptional signatures characteristic of the bipotential progenitor population and examine the nature and extent of similarities and differences between the MEP compared to its MEG and ERY progeny. Our findings confirm a high similarity in transcriptomes between MEP and MEG while, in contrast, execution of the ERY program involves active induction and repression of specific genes. Further analysis of ATAC-seq data suggest a resolution for the inconsistent grouping of MEP based on transcriptomes *versus* chromatin. A subset of the genes that are strongly up-regulated in ERY show evidence of open chromatin at their candidate *cis*-regulatory elements (cCREs) in MEP, whereas a subset of genes associated with multipotent progenitors that continue to be expressed in MEG have lower ATAC-seq signals in MEP. These patterns contribute to the clustering of MEP and MEG with other progenitors by RNA-seq but clustering of MEP with ERY by chromatin accessibility.

## Results

### RNA-seq in MEP, MEG and ERY

The levels of polyadenylated (polyA+) RNAs in ERY, MEG, and MEP (Figure 1B) were measured using directional RNA-seq. ERY and MEG were isolated from mouse fetal liver, and MEPs were isolated from adult mouse bone marrow by fluorescence activated cell sorting (FACS, Figure 1C). MEPs formed megakaryocytic and erythroid colonies almost exclusively (Figure 1D). After performing strand-specific RNA-seq (39) on two biological replicates for each sample, the reads were processed through Cufflinks and Cuffdiff (40–43) to obtain measures of expression levels (Supporting Information Figure S2) for 22,977 RefSeq genes, including genes coding for proteins as well as known non-coding RNAs in the three cell types. The replicates were highly concordant in expression levels (non-zero log2 FPKMs), with Spearman correlation coefficients of 0.97, 0.84, and 0.91 for MEP, MEG, and ERY, respectively (Figure 1E). The quality and reproducibility of the RNA-seq data are illustrated for the MEG- and ERY-expressed *Zfpm1* gene (encoding FOG1) and for lineage-specific genes, *Slc4a1* (encoding the ERY anion transporter Band 3), *Slc25a39* (encoding a mitochondrial glutathione transporter), and *Itga2b* (encoding the MEG integrin alpha 2b) (Figure 1F).

### Distributions of gene expression during erythro-megakaryopoiesis

Within each cell type, most genes were expressed at low or undetectable levels, while a minority of genes were expressed at moderate to high levels (Figure 2A). The expression levels for genes from other cell lineages were used to define the threshold for assigning a gene as expressed. Specifically, the RNA-seq fragments mapping to the lymphoid gene *Pax5* and the muscle determination gene *Myod1* gave a signal below a log2 FPKM of 3 in these deeply sequenced samples (Figure 2A), and thus we considered genes with expression levels at or below this threshold to be silent. Using this conservative, stringent threshold allowed us to focus on the robustly expressed genes in each cell type.

**Figure 2.**
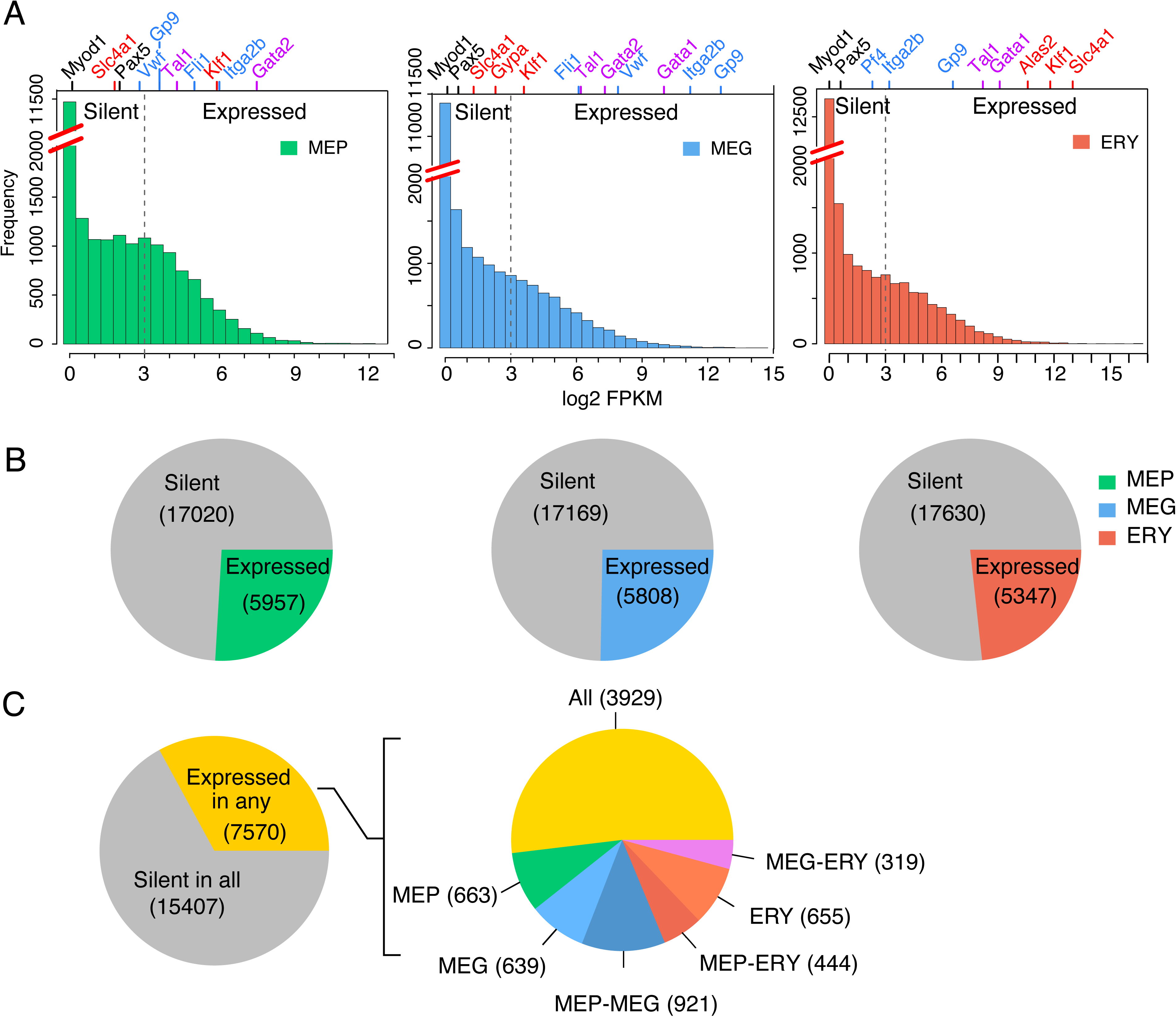
Distributions of gene expression during erythro-megakaryopoiesis. (A) Histograms showing the distribution of gene expression levels (log2 FPKM for pooled replicates plotted in bins along the X-axis) in the three cell types. Histograms are gapped along the Y-axis to better display the frequencies of moderate and highly expressed genes. Marker genes are indicated along the top X-axis, lined up at their level of expression on the bottom X-axis. Colors of gene names indicate lineage affiliation; ERY genes are in red, MEG genes in blue, genes common to both are in violet, and non-myeloid or non-hematopoietic genes are in black. The dotted line at log2 FPKM =3 indicates an empirically determined threshold for reliable detection of expression. (B) Pie charts showing the extent of transcription in each lineage. Grey indicates number of silent genes and a non-grey color (MEP = green, MEG = blue and ERY = red) indicates number of expressed genes. (C) Pie charts showing the extent of shared and lineage-specific transcription. The pie chart on the left shows the number of genes expressed in any lineage. The pie chart on the right shows the distribution of the 7,570 expressed genes across one, two, or all three lineages.

Applying this threshold, we estimate that 23 – 25% of the genes were expressed in each cell type (Figure 2B). We identified 7570 genes that were expressed in at least one of the three cell types (Figure 2C, left). About a quarter of those (1957 genes) were expressed in only one of the three cell types, divided almost equally among the three (Figure 2C, right). About half (3929 genes) were expressed in all three cell types, indicating a large sharing of the transcription program among the progenitor and both daughter lineages. Considering genes expressed in two cell types, it is notable that MEP shared a larger number of expressed genes with MEG (921) than with ERY (444). Thus, the MEP transcription program includes genes also expressed in the daughter lineages, and substantially more (χ^2^ p-value < 2.2e-16) of them are expressed in MEG than in ERY.

### MEPs are more similar transcriptionally to megakaryocytes than to erythroblasts

We compared the global transcriptomes of MEP, MEG, and ERY using three independent approaches. First, we grouped cell types by unsupervised hierarchical clustering, using correlation coefficients (Pearson’s *r*) between transcriptome profiles of all 7570 expressed genes as a distance metric. To reduce the impact of the lower correlation between MEG replicates on the overall hierarchy, we used pseudoreplicates from pooled reads of the replicates (see Methods). We found that the MEP grouped together with MEG to the exclusion of ERY (Figure 3A), showing that the transcriptome profile of MEPs was globally more similar to that of MEG than ERY.

**Figure 3.**
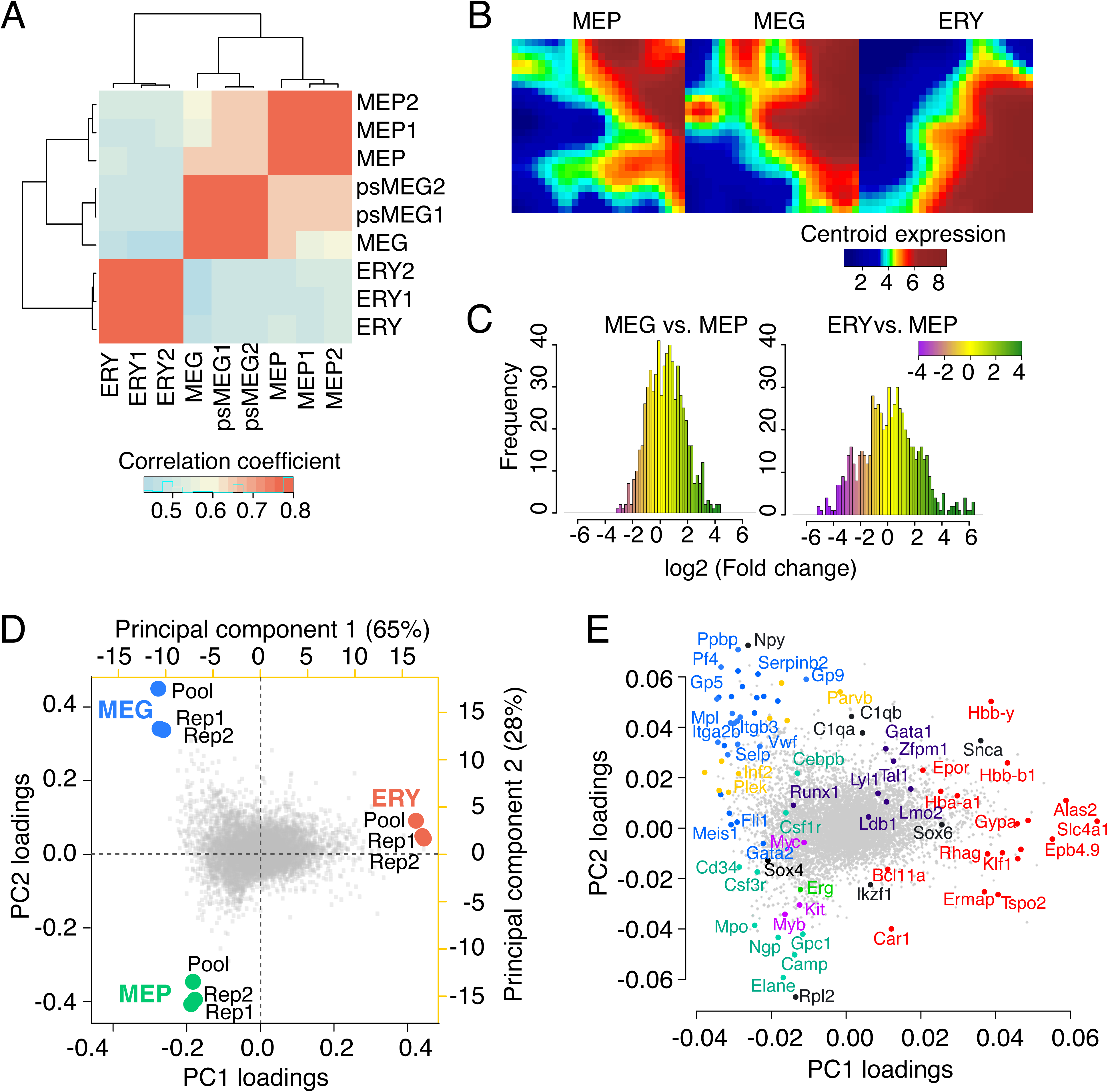
Global relationships among MEPs, MEG, and ERY transcriptomes. (A) Hierarchical clustering of replicate expression levels from MEP, MEG and ERY based on Pearson’s correlation coefficient as a distance metric. Dendrograms indicate the relationship between the cell types. Both replicate and pooled data are shown, with pseudoreplicates being used for MEG. (B) Self-organizing maps comparing expression trends across cell types generated using the GEDI tool. Groups of genes with similar expression patterns across cell types are spatially clustered together. Each tile representing a minicluster of genes is colored by its centroid expression level. (C) Histograms depicting distribution of fold changes of each tile in differential SOMs for MEG vs. MEP and ERY vs. MEP. Colors indicate level of signed fold change. (D) Lineages were clustered using principal component analysis based on replicate and pooled expression levels in each of the 7,570 expressed genes and the results were graphed as a biplot. On the bottom X-axis and left Y-axis (black), the biplot reflects the individual contribution of each gene (loadings, in gray dots) towards principal components (PCs) 1 and 2. The top X-axis and right Y-axis (in yellow) reflect the principal component (PC) scores of each sample (red, green, blue squares). PC1 separates MEP-MEG from ERY and PC2 separates MEP from MEG. (E) Informative individual genes contributing to these PC scores are labeled at the dot corresponding to the loadings on PC1 and PC2. Blue dots indicate MEG genes, yellow dots are for cytoskeletal proteins, teal dots indicate genes of non-MEG-ERY myeloid lineages, purple indicates genes for both MEG and ERY lineages and red indicates ERY genes.

We then used a machine-learning method to compare genome-wide expression profiles. Self-organizing maps (SOMs) generated with the GEDI tool (44) illustrated broad expression trends across lineages. The SOMs were built from mini-clusters of co-expressed genes whose spatial locations were preserved across maps of each cell type (45). Thus, the SOMs allow direct comparisons between expression levels of specific groups of genes and reveal the overall degree of differences between maps. Each tile in a SOM corresponds to the same group of genes across maps, and the color of the tile reflects the centroid expression level of its group of genes (Figure 3B). Tiles with similar expression levels across cell types were spatially clustered together. The patterns in the SOMs showed that the MEP and MEG were more similar to each other than either was to ERY (Figure 3B). Differential maps depicting the log of the fold-change between lineages showed this trend quantitatively (Figure 3C), with more tiles in ERY than MEG with large changes in expression when compared to MEP.

Finally, we examined the global relationships among the cell types using principal components analysis (PCA). The PCA projected our transcriptome data onto a new coordinate system of axes called principal components (PCs), which for each cell type was a linear combination of the expression level of each individual gene. Since the first few PCs can explain a large amount of variability in the data, this method reveals the most informative relationships among cell types while reducing dimensionality and retaining information on the contribution of each individual gene. The PCA results (Figure 3D) show groupings of the lineages along the first and second principal components as well as the contribution of each of the 7,570 genes (loadings) to these groupings. Replicate and pooled samples belonging to each cell type clustered faithfully for each lineage. Principal component 1 (PC 1), the axis that explained the largest amount of variance in the data (65%), separated ERY from MEG and MEP. The fact that the major component to the variance in expression for all genes separated ERY from the other two cell types shows again that MEPs are closer in expression pattern to MEG. The second principal component (PC 2) explained an additional 28% of the variance and separates MEG from MEP.

As expected, genes with high positive loadings along PC1, which separates ERY from MEP-MEG, were well-known erythroid markers (red dots, Figure 3E). Genes with negative loadings on PC1 contributed to the MEP and MEG grouping. Those with positive loadings on PC2 contributed more to the MEG lineage than to MEP, and they included well-known megakaryocytic genes. Strikingly, several genes contributing to the MEP lineage (i.e., with negative values along PC1 and PC2) were also highly expressed other myeloid cells, such as granulocytes and macrophages. Examples include genes encoding myeloperoxidase (*Mpo*), neutrophil elastase (*Elane*), and neutrophil granule protein (*Ngp*). We conclude from the results of three independent computational approaches analyzing the polyA+ RNA that the transcriptome of MEPs was more closely related to MEG than ERY.

### Erythroid transcription program undergoes a greater degree of upregulation

Since the MEP transcriptome preferentially expresses MEG genes relative to ERY genes, we hypothesized that erythroid genes would need to be activated in ERY relative to their expression in MEP. We tested this hypothesis by examining the changes in gene expression after MEPs differentiate into MEG and ERY. To ensure that our results are robust, we used two different methods to characterize differentially expressed genes, Cuffdiff (41, 42) and k-means clustering, evaluating the hypothesis in both sets of results as well as a consensus set.

First, pairwise differential expression tests using Cuffdiff compared each differentiated cell type to the MEP as a reference. Genes were declared significantly changing if they met both of two criteria: (i) their differential expression passed an FDR (false discovery rate) threshold of 0.05, and (ii) they were expressed in at least one of the two lineages being compared. Based on the direction of change, genes were categorized as up-regulated (U), down-regulated (D), or not changing significantly (N) in the differentiated cells as compared to the progenitor (Figure 4A,B). The up-regulated category includes both genes activated from a “silent” state in MEP to “expressed” in the daughter cells and genes expressed at a low level MEP but induced to a higher level in the daughter cells. A composite notation conveyed differential expression in both lineages, e.g. “UN” denotes a gene up-regulated in MEG and not changing significantly in ERY, as compared to MEP. This process assigned 6,331 differentially expressed genes to nine such expression-change categories.

**Figure 4.**
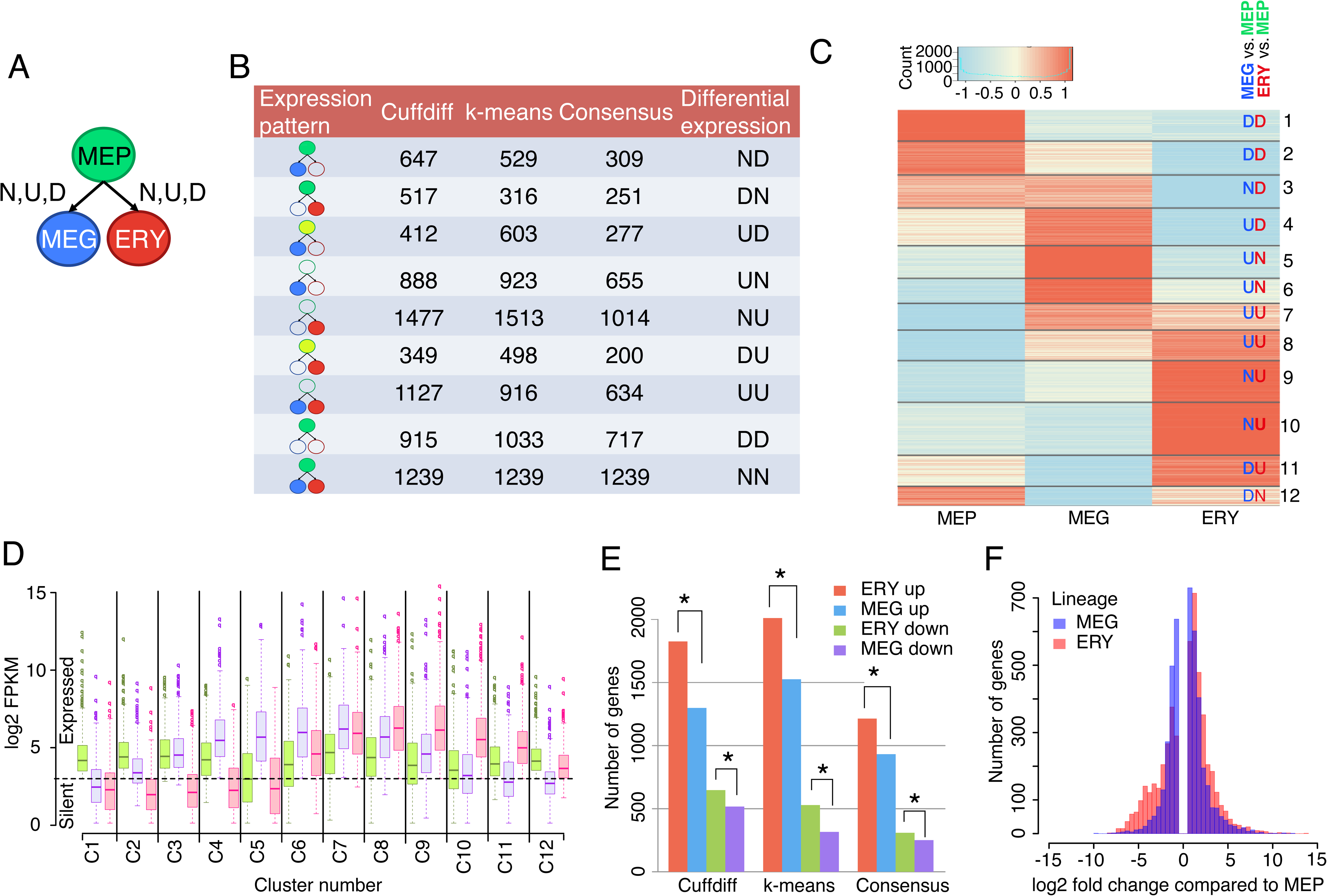
Erythroid program is characterized by a greater degree of induction. (A) Schematic showing the pairwise differential expression tests performed for MEG (blue oval) and ERY (red oval) using MEP (green oval) as a reference. Possible outcomes are induction (“U”), repression (“D”) and no change (“N”). (B) Table showing the number of genes declared differentially expressed using each method (Cuffdiff and k-means) and their consensus. The cartoons indicate expression patterns: red, blue or green filled ovals indicate expression in ERY, MEG and MEP respectively. Absence of a color (empty ovals) indicate no expression in a lineage. (C) Heatmap depicting genes clustered based on expression level in each of the three cell types. Red = relatively higher expression and blue = low expression, as represented by row-standardized log2 FPKMs. (D) Unstandardized log2 FPKMs are plotted for each sample in each cluster, to assign each cluster to an expression category. (E) Barplots summarizing the number of induced and repressed genes in ERY and MEG, as compared to MEP. Gene categories are “ERY up” (DU + NU), MEG up (UD + UN), ERY down (ND) and MEG down (DN). Stars indicate significant differences between categories indicated, using the Chi-sq test and Bonferroni-corrected p-values are reported. (F) Histograms showing the fold change for differentially expressed genes in ERY and MEG, as compared to MEP.

We then used k-means clustering to classify the differentially expressed genes into categories. This approach differs from the pairwise comparisons in Cuffdiff because it considers expression levels across all three cell types simultaneously. Varying k, the number of clusters, from 1 to 25 showed that at k = 12, the within cluster sum of squares was minimized (Supporting Information Figure S3), and hence the genes were tightly clustered. The 12 clusters grouped informative subsets of genes by expression pattern across the three cell types, revealing both lineage-specific and lineage-shared genes (Figure 4C, D).

Despite showing similar trends for the categories of differentially expressed genes, Cuffdiff and k-means clustering placed different numbers of genes in each category (Figure 4B). Therefore, we generated a third, consensus set of 4057 differentially expressed genes, limiting the genes in each category to the ones that were assigned to the same category by both methods (Figure 4B). The expressed genes, their log2 FPKM in each cell type, and their differential expression cluster assignments are provided in Supporting Information Table 1.

**Table 1.** Enumeration of cCRE-gene pairs in categories of ATAC-seq patterns across cell types for enrichment analysis.

| A. Counts of all cCRE-gene pairs in categories |  |  |  | B. Enrichment or depletion in informative categories |  |  |  |  |  |
| --- | --- | --- | --- | --- | --- | --- | --- | --- | --- |
| Cell type |  |  |  | B1. ATAC-seq higher in MEP and ERY than MEG |  |  |  |  |  |
| MEP | MEG | ERY | Number | Cell Type |  |  | Number of cCRE-gene pairs |  |  |
|  |  |  |  | MEP | MEG | ERY | All genes | Cluster 10 | Cluster 3 |
| H | H | H | 3418 |  |  |  |  |  |  |
| H | H | L | 37 | H | L | H | 187 | 50 | 1 |
| H | H | M | 669 | H | L | M | 246 | 36 | 14 |
| H | L | H | 187 | H | M | H | 632 | 159 | 9 |
| H | L | L | 98 | M | L | H | 541 | 160 | 9 |
| H | L | M | 246 | M | L | M | 3370 | 645 | 118 |
| H | M | H | 632 | Number pairs with higher signal in MEP and ERY |  |  | 4976 | 1050 | 151 |
| H | M | L | 90 | Total pairs in all genes or clusters |  |  | 48701 | 5223 | 4015 |
| H | M | M | 677 |  |  | percent | 10.2 | 20.1 | 3.8 |
| L | H | L | 532 |  |  | odds ratio |  | 2.21 | 0.34 |
| L | H | M | 43 |  |  | 95% CI |  | 2.05-2.38 | 0.29-0.41 |
| L | L | H | 25 |  |  |  |  | Enriched | Depleted |
| L | L | L | 4549 |  |  |  |  |  |  |
| L | L | M | 730 | B2. ATAC-seq low in MEP but medium or high in MEG |  |  |  |  |  |
| L | M | H | 3 | MEP | MEG | ERY | All genes | Cluster 10 | Cluster 3 |
| L | M | L | 8482 | L | H | H | 0 | 0 | 0 |
| L | M | M | 1051 | L | H | L | 532 | 19 | 71 |
| M | H | H | 1100 | L | H | M | 43 | 7 | 2 |
| M | H | L | 456 | L | M | H | 3 | 0 | 0 |
| M | H | M | 929 | L | M | L | 8482 | 582 | 941 |
| M | L | H | 541 | L | M | M | 1051 | 132 | 50 |
| M | L | L | 2825 | Number pairs L in MEP, H or M in MEG |  |  | 10111 | 740 | 1064 |
| M | L | M | 3370 | Total pairs in all genes or clusters |  |  | 48701 | 5223 | 4015 |
| M | M | H | 1366 |  |  | percent | 20.8 | 14.2 | 26.5 |
| M | M | L | 4969 |  |  | odds ratio |  | 0.63 | 1.38 |
| M | M | M | 11676 |  |  | 95% CI |  | 0.58-0.68 | 1.28-1.48 |
|  |  | total | 48701 |  |  |  |  | Depleted | Enriched |

All three sets of differentially expressed genes strongly support the hypothesis that induction is a dominant mode of regulation in erythroid differentiation. Considering the genes with expression changes in the MEG and ERY lineages, the number induced when MEP differentiate into ERY is consistently higher than the number induced when MEP follow the alternate fate to MEG (Figure 4E) in all three sets. Also, the number of genes repressed as MEPs differentiate to ERY was greater (Figure 4E, Bonferroni-adjusted χ^2^ p-values < 0.05 for all sets) than the number repressed in MEP. Furthermore, a larger number of genes showed higher and lower levels of expression change during differentiation to ERY than to MEG (Figure 4F, Kolmogorov-Smirnov (KS) test p-value < 2.2e-16). These results show that a larger number of genes are actively regulated, mainly by induction, in erythropoiesis than in megakaryopoiesis after the MEP stage.

### Enrichment of function-related terms in lineage-specific and shared genes

We used the GREAT computational tool (46) to investigate more completely which functions were enriched in the genes in the nine expression categories (Figure 5A). As expected, the annotations of genes upregulated specifically in ERY, i.e. expression categories NU and DU, were enriched in terms related to erythroid functions such as erythrocyte morphology and heme biosynthesis. The ERY-specific genes were also enriched in terms associated with cell growth and proliferation, consistent with the rapid cell division that occurs in early erythroid differentiation. Likewise, the annotations of genes upregulated specifically in MEG, i.e., expression categories UN and UD, were enriched in terms related to megakaryocyte functions such as platelet activation and blood coagulation (Figure 5A).

**Figure 5.**
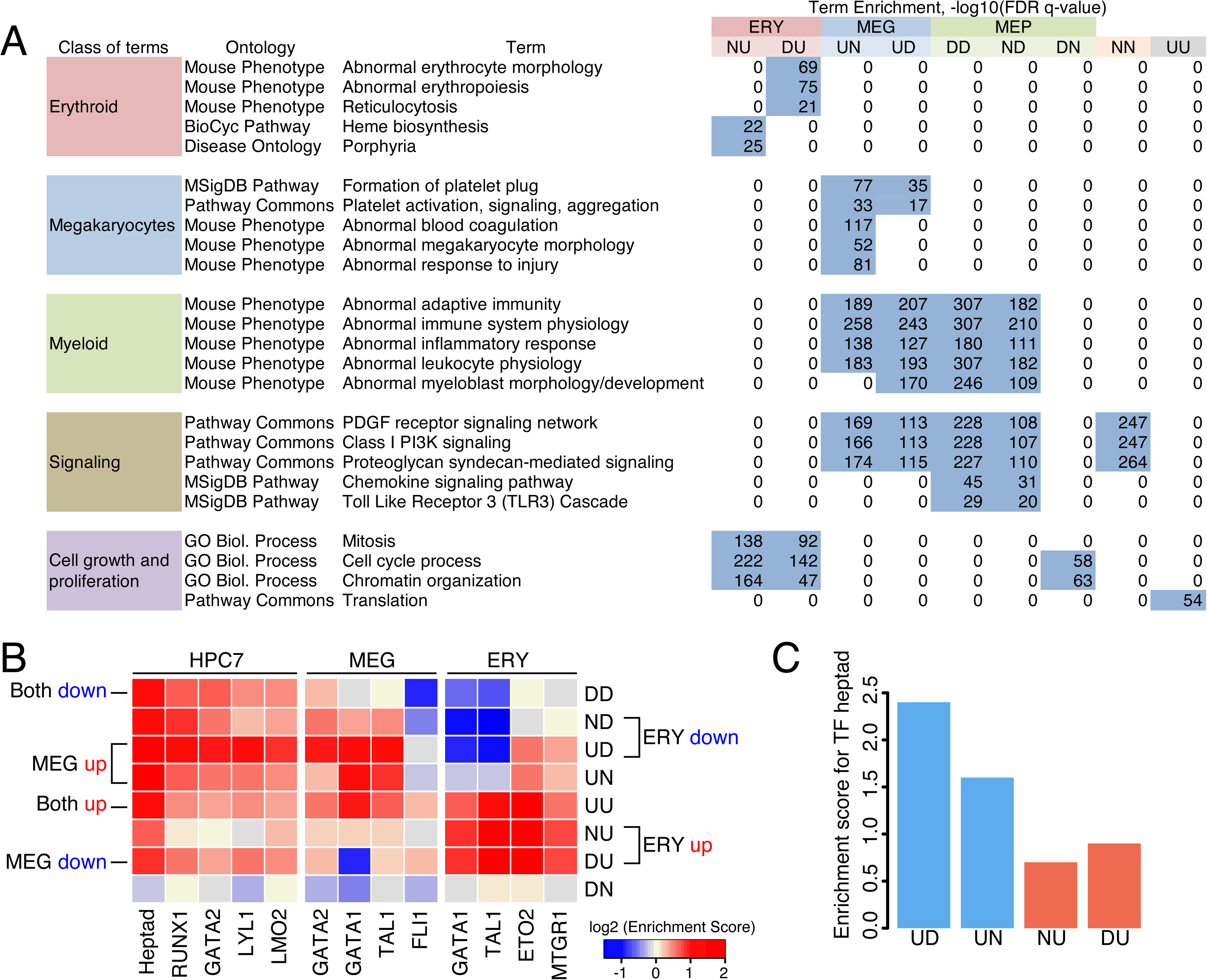
Functional term enrichments and transcription factor occupancy for expression categories. (A) Function-related terms that are significantly enriched (binomial and hyper-geometric FDR < 0.05) in the gene sets for each differential expression category were determined across multiple ontologies using GREAT (46). Examination of the 50 most significant terms for each expression category revealed frequent terms in five classes of terms, which are shown along with illustrative specific terms and the negative logarithm of the FDR q-values in each expression category, with non-zero values highlighted. (B) Heatmap showing the enrichment of TF occupancy (columns) in genes in each expression category (row). TF peaks were assigned to a gene if they were found in the gene neighborhood, defined as the interval from 10 kb upstream of the transcription start site (TSS) to 10 kb past the polyA addition site^38^. Each gene can thus be assigned multiple peaks and peaks could also be assigned to multiple genes. Fold enrichment (or depletion) was computed as the percentage of genes in an expression category occupied by each TF divided by the percentage occupied in a background set and expressed as the log_2_-fold enrichment (represented by the color of each tile). (C) Barplot showing enrichment of occupancy of the TF heptad in various expression categories.

We were particularly interested in the functions of the genes expressed in MEP but that were down-regulated in both MEG and ERY, i.e. category DD. The annotations for these genes were highly enriched in terms associated with other myeloid cells, e.g. adaptive immunity and inflammatory response, as well as several signaling pathways, such as PDGF receptor signaling and PI3K signaling (Figure 5A). Notably, these term enrichments were also observed for genes expressed in MEG, both those that were up-regulated during differentiation to MEG (UN and UD categories) as well as those that were expressed in MEP and retained in MEG (ND category). We conclude that the expression profile of the MEP population contains genes characteristic of myeloid cells, including those encoding several signaling pathways. Expression of some of these are retained in MEG, but they are not expressed in ERY.

### Transcription factor occupancy in differentially expressed genes

The analysis of differentially expressed genes showed that a substantial number were co-expressed in MEP and MEG whereas another substantial set was induced specifically in ERY. We hypothesized that differences in TF occupancy could explain, at least in part, these distinctive expression patterns, and we investigated the predicted association by calculating enrichment of the genes for chromatin occupancy by key TFs in relevant cell types. Specifically, we used ChIP-seq peak calls for TF occupancy in a cell line model for multipotent hematopoietic progenitor cells HPC7 (47), in MEG (19), and in erythroid cells (19, 48). (Peak statistics and sources are in Supporting Information, Figure S4.) The data from HPC7 cells was used as a proxy for MEPs since the latter could not be obtained in sufficient numbers to perform standard ChIP-seq assays. Enrichment or depletion of occupancy by each TF for genes in each expression category was computed based on the numbers of TF occupancy peaks observed in each gene neighborhood (the entire gene with extension of 10kb on each end) compared to the numbers of peaks in the set of 1239 genes with no change in expression. This analysis considers numbers of occupied sites in the genes, but it does not investigate or compare which regulatory elements in the genes are bound. Some elements could be bound by different sets of TFs in different cell types.

Genes in almost all the differential expression categories were enriched for occupancy by TFs in the progenitor cell line model HPC7 (Figure 5B). The exception were genes in DN category, which presented a distinctly different pattern of depletion or very low enrichment for TFs across all cell types. Genes upregulated in both lineages, category UU, were further enriched for occupancy by TFs in both ERY and MEG beyond the more modest occupancy in HPC-7. The induction of genes in the ERY and MEG lineages showed distinctive patterns. Genes induced specifically in ERY (categories NU and DU) were strongly enriched for occupancy by the examined TFs in erythroid cells but modest to no enrichment for TFs in HPC7 and MEG cells. This result indicates new binding by TFs at these genes during ERY differentiation. In contrast, genes induced specifically in MEG (categories UD and UN) were strongly enriched for most TF occupancy in *both* megakaryocytes and hematopoietic progenitor cells. While occupancy by the TF heptad (47) was enriched in several expression categories, we observed a greater degree of enrichment of occupancy by the TF heptad for induction in MEG than in ERY (Figure 5C). Using HPC7 data as a proxy for MEP, we infer from these results that many genes up-regulated in MEG likely were already bound by TFs in MEP.

Genes downregulated in both lineages (category DD, Figure 5B) and those repressed in the ERY lineage (categories ND and UD) were enriched for TF occupancy in the hematopoietic progenitor cell line, but they were reduced for occupancy by many TFs in the erythroid and megakaryocyte cells. A similar trend was found for the DU category of genes repressed specifically in MEG, which was depleted for occupancy by GATA1 and had lower enrichment for other factors in MEG compared to HPC7. These patterns are consistent with activation of the genes by TF occupancy in HPC7, but the lineage specific TFs in MEG did not replace those progenitor-bound factors, likely leading to repression during differentiation. The GATA and related factors provide striking examples of this TF loss. Binding by GATA2 in HPC7 cells was enriched in genes in the categories that were downregulated in at least one lineage (categories DD, ND, and DU, not including DN), but binding by the paralogous GATA1 and the associated factor TAL1 was depleted upon repression in erythroid cells (DD and ND). Similarly, the occupancy for GATA1, TAL1, and FLI1 was less enriched or was depleted in MEG for repressed genes (categories DD and DU).

Taken together, these trends show that induction of genes in ERY frequently involved new binding of TFs, induction of genes in MEG occurred with retention or enhancement of TF binding compared to the pattern in HPC7 cells, and down-regulation in both lineages was frequently accompanied by, and likely resulted from, loss of occupancy by key TFs.

### Discordance between chromatin- and RNA-based distances for MEP

The MEP population was grouped with other blood cell types differently depending on whether RNA or chromatin features were examined, with MEP clustering with multipotential progenitors and MEG by RNA distances but with ERY by chromatin accessibility measures (Figure 1A). In many studies, chromatin accessibility and modifications such as H3K4 methylation have been strongly associated with RNA levels, and this pattern was observed for many genes in our work. For example, the gene *Slc25a4* is expressed in MEPs and MEGs but not erythroid cells, and ATAC-seq peaks were observed at the TSS and upstream of the gene only in the cell types expressing this gene (Figure 6A).

**Figure 6.**
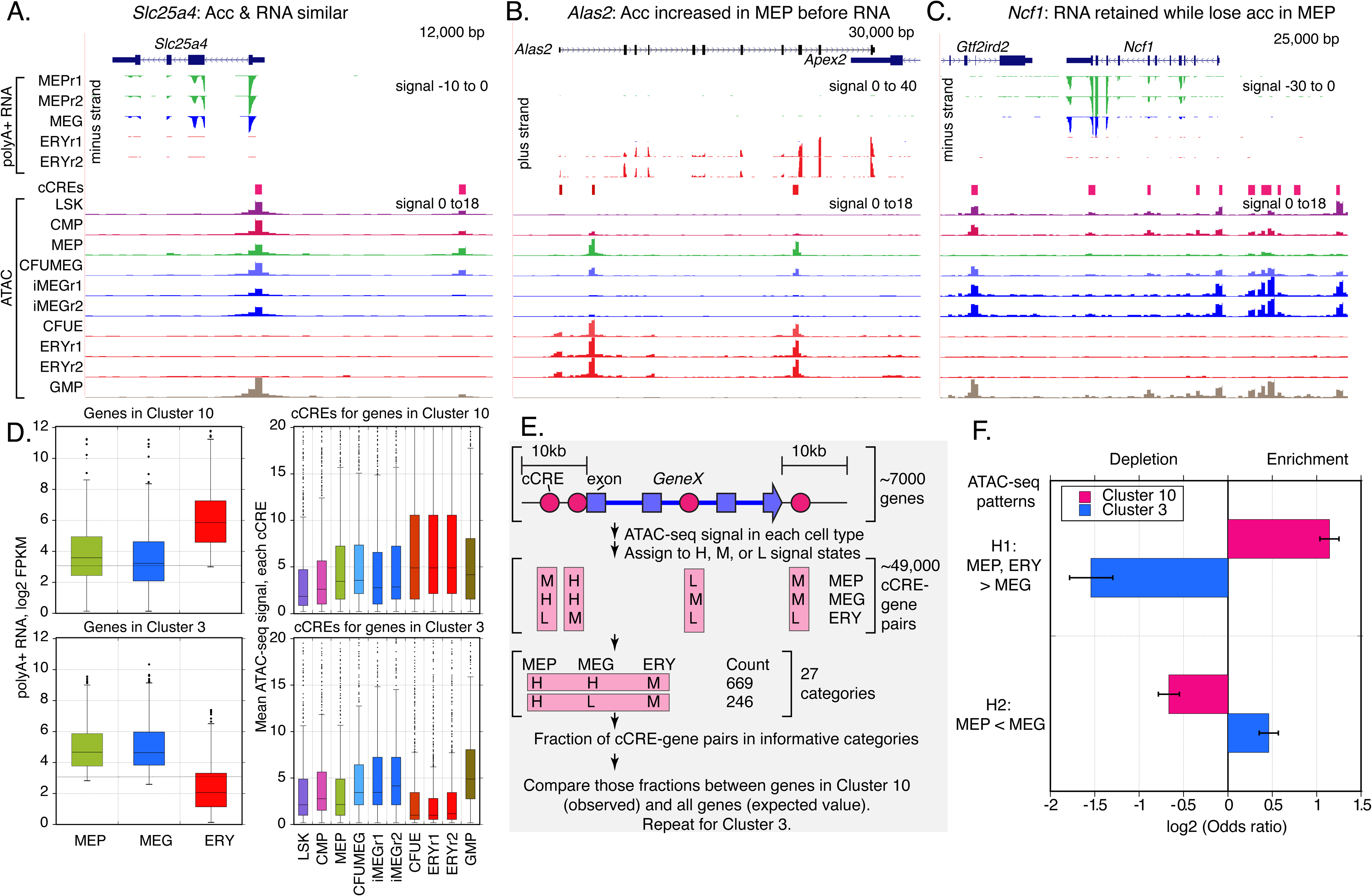
Examples and patterns in genes with discordant chromatin accessibility and RNA patterns across MEP, MEG, and ERY. (A) – (C) Examples of genes showing either (A) concordance of RNA-seq and ATAC-seq patterns, (B) precocious actuation of cCREs inferred from ATAC-seq compared to RNA-seq patterns, or (C) high levels of RNA in MEP despite a low ATAC-seq signal in cCREs. For each panel, the gene model from Gencode VM23 on the mm10 mouse genome assembly is shown at the top, followed by tracks of stranded RNA-seq data, showing only the data on the transcribed strand (synonymous with the RNA). The y-axis for the signal is the same for all tracks in a panel; negative numbers are used for signal on the minus strand. The cCREs determined from normalized ATAC-seq signals across mouse blood cell types from the VISION project (24) are shown above tracks of ATAC-seq data for cell types in the differentiation series from the stem and progenitor population LSK, through the common myeloid progenitor (CMP), MEP, precursors to MEG (CFUMEG), immature MEG (iMEG), precursors to ERY (CFUE), and ERY, and also including the granulocyte monocyte progenitor population (GMP) for comparison. The ATAC-seq signal normalization was computed in 200bp bins across the genome, and thus the signal (negative log base 10 of the p-value for difference from expectation of a binomial distribution) is shown at a resolution of 200bp. The same y-axis limits were set for signal across cell types. Additional abbreviations are Acc for chromatin accessibility; r1 and r2 for replicates 1 and 2. (D) Distributions of RNA-seq signals for genes and ATAC-seq signals for cCREs associated with those genes in differential expression Clusters 10 and 3. The distributions are summarized as box plots with the box extending from the first to the third quartile, the line indicating the median, and the whiskers extending to the most extreme data point that is no more than 1.5 times the interquartile range from the edge of the box, and with outliers shown as points. (E) Steps in compiling and categorizing cCRE-gene pairs for assessment of frequency of fit to hypothesized patterns to explain discordance in grouping of MEP with other cell types. (F) Assessment of the frequency of ATAC-seq patterns predicted by two hypotheses (H1 and H2) to explain discordance in grouping of MEP with other cell types, comparing frequency of the patterns as observed for cCRE-gene pairs in Clusters 10 and 3 with those for all gene genes as expected. The evaluation used Fisher’s Exact Test, with results expressed as an odds ratio, which was plotted as the log base 2 to distinguish enrichment (positive values) from depletion (negative values). The error bars cover the 95% confidence interval for the odds ratios. All comparisons were significant by Fisher’s Exact Test (p<0.0000).

We reasoned that the discordance in cell type grouping could arise from sets of genes that showed a pattern of chromatin accessibility that differed from the RNA pattern across the three cell lineages. We searched for such examples in potentially informative clusters of genes defined by the robust differential expression assignments from the polyA+ RNA data, comparing the RNA levels with the normalized ATAC-seq signals in blood cell types for cCREs from the VISION project (23, 24). One hypothesis to explain the greater similarity of MEP and ERY in chromatin accessibility compared to RNA was precocious actuation (accessibility to nucleases in chromatin) in MEP of cCREs for genes that would be expressed primarily in ERY. This hypothesis predicts that chromatin accessibility would be observed in the MEP population for genes with RNA levels that were low or undetectable in MEP but high in ERY. The genes in differential expression Cluster 10 (Figure 4C, D) were candidates for manifesting the predicted pattern because they had low expression in MEP and MEG but high in ERY. Indeed, the *Alas2* gene fits with the predicted pattern of abundant RNA in ERY but not in MEP or MEG, coupled with unexpected strong ATAC-seq peaks at cCREs in MEP, along with the expected ones in erythroid cells (Figure 6B).

A second hypothesis to explain the discordance was that some genes expressed in MEP and MEG had low or undetectable ATAC-seq signal at their associated cCREs in MEP, thereby leading MEP to be grouped with MEG by RNA data but not by chromatin accessibility. The genes in differential expression Cluster 3 (Figure 4) were expressed in MEP and MEG but not in ERY, and therefore they were candidates for manifesting the pattern predicted by the second hypothesis. The gene *Ncf1* conforms to the predicted pattern, with high RNA-seq signals in MEP and MEG (not ERY) but low signal for chromatin accessibility in MEP along with high signal in MEG (Figure 6C). Some of the genes with the discordance between levels of RNA and chromatin accessibility in MEP were in active chromatin in earlier multi-lineage progenitor cells, as illustrated by the ATAC-seq signals in cCREs for *Ncf1* in LSK, CMP, and GMP (Figure 6C). Possibly these genes that were expressed in earlier progenitors, MEP, and MEG have lost chromatin accessibility in some of the sub-populations of MEP, perhaps those that are more likely to differentiate into erythroid cells, while retaining substantial stable polyA+ RNA.

We compared the patterns of RNA and chromatin accessibility more completely by collecting the sets of cCREs associated with each gene expressed in at least one of MEP, MEG, or ERY and tabulating the mean normalized ATAC-seq signal in each of these three cell types using information in dbCRE from the VISION project (24). The cCREs found within the extended gene body were considered associated with a gene (Figure 6E, see Methods). The distributions of ATAC-seq signals in cCREs associated with genes in Cluster 10 indicate some potential support for the hypothesis of precocious actuation of cCREs in MEP, with a trend toward higher ATAC-seq signals in MEP than in MEG (Figure 6D). The distribution of ATAC-seq signals for cCREs associated with genes in Cluster 3 indicate support for the second hypothesis, with a substantially lower distribution in MEP than in MEG (Figure 6D). However, many different patterns of chromatin accessibility could be contributing to the broad distributions of signals observed.

We then adopted a categorization strategy to summarize an informative and comprehensive set of patterns for the ATAC-seq signals in the three cell types. Three ATAC-seq signal states, high (H), medium (M), and low (L), were defined using thresholds derived from the distributions of normalized ATAC-seq signals in the mouse blood cell types (see Methods). Each of the 48,701 cCRE-gene pairs for the 7,590 expressed genes was assigned to one of 27 possible categories of ATAC-seq signal state across the three cell types (Table 1A, Figure 6E). This enumeration of cCRE-gene pairs in the signal states was used to test predictions of the two hypotheses to explain the discordance in cell type grouping by RNA versus chromatin accessibility. Each hypothesis predicted that cCRE-gene pairs would be in a particular subset of the chromatin state categories. The proportion of cCRE-gene pairs for all expressed genes that were assigned to that subset of categories served as the expected value, and the proportion of cCRE-gene pairs for genes in informative differential expression Cluster 10 and in Cluster 3 served as the observed values (Table 1B; Figure 6E). Specifically, we evaluated whether cCREs associated with genes in Cluster 10 (high expression only in ERY) were enriched for cCREs precociously actuated in MEP (hypothesis 1) by calculating the proportion of cCRE-gene pairs in signal state categories with ATAC-seq signals higher in MEP and ERY than in MEG. These cCREs for genes in Cluster 10 were significantly enriched (odds ratio = 2.21) while the cCREs for genes in Cluster 3 (high expression in MEP and MEG, not ERY) were significantly depleted (odds ratio = 0.34; Table 1B1; Figure 6F). As a test of the second hypothesis, we evaluated whether cCREs associated with genes in Cluster 3 were enriched for the predicted ATAC-seq patterns with lower signal in MEP than in MEG. The cCREs associated with Cluster 3 genes were significantly enriched for an ATAC-seq signal higher in MEG than in MEP (odds ratio = 1.38), whereas the cCREs associated with Cluster 10 genes were significantly depleted (odds ratio = 0.63; Table 1B2; Figure 6F). All four comparisons were significant (p<0.000) by Fisher’s exact test.

Based on these analyses, we conclude that both (a) precocious actuation in MEP of cCREs for erythroid genes and (b) persistence of stable RNAs for some other genes in MEP despite a loss of chromatin accessibility contribute to the discordance in the grouping of MEP with other blood cell types depending on the experimental modality used to generate the distance metrics.

## Discussion

MEP comprise a population of cells, derived from the common myeloid progenitor (CMP) cell population, that is capable of differentiating into MEG and ERY. The MEP do not represent a static, stable stage but rather they are a prominent intermediate population in the progressive differentiation from LSK through CMP and MEP to MEG and ERY. MEP are not a homogeneous cell population, and human MEP can be resolved into at least three subpopulations. The RNA-seq and ATAC-seq data presented and examined in this paper provided sensitive measurements, based on deep sequencing, of the composite transcriptomes and chromatin accessibility landscapes of the MEP population and of the more homogeneous ERY and MEG cell preparations. Examples of maintenance, repression, and induction of genes were found in both the lineages from MEP to either MEG or ERY. However, despite the confounding effects of heterogeneous subpopulations in MEP, we observed two distinct modes that dominate regulation in the two lineages.

Commitment to megakaryopoiesis features maintenance of a MEG program largely present in the MEP population, whereas erythroid differentiation requires a larger role of active induction and repression, i.e. a re-wiring of the regulatory program. The MEG and MEP transcriptomes were highly similar, as revealed by the shared expression of over twice as many genes between MEP and MEG than between MEP and ERY, the grouping of MEP and MEG to the exclusion of ERY in hierarchical clustering and self-organizing maps, and the observation by PCA that the greatest variation in transcriptomes separated ERY from MEP and MEG. Analysis of differentially expressed genes revealed that differentiation to ERY involved the induction and repression of more genes than was found for differentiation to MEG. These conclusions about different modes of regulation in the two lineages were supported by analysis of transcription factor occupancy, which showed that the genes differentially expressed genes on the lineage to MEG were already enriched for occupancy of key transcription factors in a cell line model for multipotent hematopoietic progenitor cells (HPCs), whereas genes induced or repressed in the lineage to ERY showed large increases or decreases in enrichment, respectively, for induction and repression.

Our conclusions on the different modes of regulation strongly support earlier reports of maintenance of TF occupancy from HPCs to MEG in contrast to loss of occupancy at repressed genes and *de novo* occupancy at induced genes in ERY (19). Furthermore, analysis of transcriptomes and chromatin accessibility across diverse multipotent and maturing blood cell types in mouse also showed substantial retention of transcripts and actuated cCREs from LSK through CMP to immature MEG, whereas significantly fewer progenitor-expressed transcripts and actuated cCREs were retained in ERY (22). The distinct programs of regulation in the MEG and ERY lineages were also supported by the observation that most genomic sites occupied by GATA1 and TAL1 differed between MEG and ERY (17, 19, 20). These different modes of regulation in the two lineages also include differences in DNA methylation, with substantial *de novo* DNA methylation during differentiation to MEG (22) in contrast to global demethylation of DNA during differentiation and maturation of ERY (22, 49). Despite the strong similarity in transcriptomes between MEP and MEG and the need for substantial regulatory rewiring in the ERY lineage, the high proliferative potential of the cells differentiating toward ERY leads to a substantial component of the MEP population committing to erythropoiesis, at least in humans (34).

Our data support some proposed processes in cellular differentiation. Several models of differentiation posit that cells with multilineage potential express many of the genes characteristic of the mature lineages, and specialization of cellular identity is achieved by progressive restriction of transcription (50–52). The broad transcription program observed in MEP and the repression of at least 1200 genes in the MEG and ERY lineages are consistent with this model. Among these MEP-expressed genes that were subsequently repressed were genes whose expression is characteristic of cells in other lineages, such as granulocytes and monocytes. Expression of those genes in MEP could reflect a cellular “memory” of greater potentiality at an earlier stage, such as CMP. Another process frequently observed is lineage priming (51), in which genes that are highly expressed in maturing lineages are also expressed at lower levels in the multipotent progenitor cells. For example, the MEP-expressed genes that continue expression in MEG but are repressed in ERY were highly enriched for MEG functions. Expression of many of these genes increased from MEP to MEG (such as cluster 4), thereby showing evidence for lineage priming. We also observed evidence of priming via chromatin accessibility, such as the genes in cluster 10 that were expressed only in ERY but which showed actuation of cCREs in MEP.

We conclude that MEPs have a strong bias, leaning toward a megakaryocyte fate at the expense of erythropoiesis. This MEG bias in MEP and reports that early hematopoietic stem and progenitor cells are biased towards a megakaryocyte fate fit with the molecular connections among platelets, hematopoietic stem cells, and endothelial cells, including similarity of cytokines or chemokine receptors expressed and the shared expression of transcription factors (53). However, differentiation of MEP to MEG is not a default program because the genome in differentiating MEG acquires substantial *de novo* methylation, which adds another layer of regulation during commitment and maturation to MEG (22).

Chromatin accessibility and histone modifications characteristic of active transcription or regulation, such as H3K4 methylation and H3K27 acetylation, are strongly associated with gene transcription. Indeed, these biochemical changes represent steps in the activation or memory of gene expression. However, multiple studies including ours have shown that the clustering of MEP with other hematopoietic cell types differs depending on the distance metric used, chromatin accessibility *versus* stable RNA accumulation. This unexpected discrepancy suggested that at least some sets of differentially expressed genes were showing an apparent dissociation between the acquisition of chromatin accessibility and activation of gene expression during differentiation of the MEP population to ERY or MEG. Our comparative analysis of chromatin accessibility and differential expression revealed two different categories of discordance between chromatin accessibility and gene expression. In the first category, regulatory elements were precociously actuated in MEP in genes that were silent in MEP but were highly expressed in ERY. The other category consisted of genes highly expressed in both MEP and MEG that showed little to no chromatin accessibility at their cCREs in MEP. These genes could include those that were expressed in earlier multipotent progenitor cells such as CMP, but they were no longer actively transcribed in MEP, reflected in a loss of chromatin accessibility, while their RNA was stable and hence still present in MEP. Genes in both categories would contribute to clustering of MEP with ERY by chromatin accessibility but clustering of MEP with MEG by RNA.

An issue for future study is to identify the molecular or cellular processes that lead to these discordances. One hypothesis is that such discordance is more likely to be observed during differentiation from progenitor cell populations, like MEP, with different subpopulations. For instance, the first category of apparently discordant genes might have cCREs highly accessible in a subpopulation of MEPs along with modest expression levels. In the heterogeneous MEP, this subpopulation could contribute measurably to the chromatin accessibility profiles but not to the RNA pool. This subpopulation may be biased toward commitment to the ERY lineage, which also would lead to substantial proliferation, which in turn would generate a high expression level of these genes in ERY. The second category of apparently discordant genes could reflect a high level of stable RNA in a subpopulation of MEP, perhaps from prior expression followed chromatin inaccessibility, or perhaps the genes continue expression in a low abundance subpopulation that did not contribute measurably to the chromatin accessibility profile. This general hypothesis of the role of subpopulations, illustrated by these scenarios, would predict that further refinement of robust subpopulations of MEP and increased sensitivity in multimodal single cell analyses of these differentiation pathways would reveal stronger concordance in the chromatin accessibility and transcriptome profiles within subpopulations.

## Experimental procedures

### Cell isolation

Primary erythroblasts and megakaryocytes were obtained from E14.5 CD-1 mouse fetal livers. ERY were isolated using immunoselection with TER119 antibodies, and MEG were obtained by culturing KIT-positive cells for 12 days in a megakaryocyte expansion medium containing thrombopoietin followed by immunomagnetic sorting, as described previously (19). Adult mouse bone marrow were FACS-sorted, selecting for Lin- (specifically Ter119-, CD11b-, Gr1-, IL7r1-, CD4-, CD8-, B220-), KIT+, SCA1-, CD34^low^, CD16/32-cells, to obtain MEPs (37).

### Colony assays for MEPs

Mouse CFU-GM, CFU-mix and BFU-E were assayed using Methocult, from Stem Cell Technologies (Vancouver; cat # 3434). Mouse CFU-MKs were assayed using Megacult, from Stem Cell Technologies (Vancouver; cat # 04974).

### RNA sequencing and read mapping

Total RNA was extracted from two independent biological replicates (5-10 million cells each) from of each type using Qiagen RNeasy kits, followed by OligodT magnetic bead selection for polyA+ RNA. Strand-selective cDNA libraries were prepared using dUTP in place of dTTP in second strand cDNA synthesis to allow its subsequent, selective digestion (39). Libraries were sequenced on the Illumina HiSeq 2000 to a minimum depth of 100 million reads pairs per sample. Reads were mapped to the mouse mm9 reference genome using the spliced read aligner TopHat2 (54, 55), which was supplied with Illumina’s iGenomes mm9 RefSeq GTF as a gene model reference. A two-step mapping strategy (56) was used to obtain and combine splice junctions from all samples that could be used to annotate mapped reads. The first round of mapping identified all novel splice junctions in each replicate, which were then combined across samples to obtain a master set of combined novel junctions. A second round of mapping was performed, with the master set of junctions supplied using option “-j” (--raw-juncs). At this step, “--no-novel-juncs” was enabled so that all mapped reads were annotated with splice junctions only from this master set, thereby obtaining assembly based on an all-inclusive set of splice junctions derived from all the samples. The sorted BAM output from the second round of mapping was used as input to Cufflinks (40–42) for quantification of expression levels for individual replicates and to Cuffdiff (43) for differential expression testing. The coverage across exons was calculated and signal tracks were generated from this BAM output using SAMTools (57), BEDTools (58), the UCSC Table Browser (59), and other UCSC utilities (60–63).

### Quantification of expression levels

We used a custom gene model annotation file in which each gene was represented by a single canonical transcript to estimate expression levels for RefSeq genes. Starting with an Illumina iGenomes RefSeq mm9 GTF, we obtained canonical transcripts for each gene from the “knownCanonical” table using the UCSC Table Browser (59) to represent that gene. For genes without any record of a canonical transcript, we chose a representative transcript based on transcript length (longest), CDS length (longest) and number of exons (greater). We excluded genes positioned on chrN_random or chrUn_random and snoRNAs matching the pattern “Snora”. This resulted in 22,977 genes, each with a single transcript. We used Cuffdiff (40–42) to identify differentially expressed genes, using this custom GTF with the parameters --dispersion-method set to per-condition, --library-type set to fr-firststrand, --max-bundle-frags = 20000000, -- min-reps-for-js-test = 2, -b for bias correction and –M to mask globin transcripts. However, regions on mouse chr11 and chr7 containing alpha-globin and beta-globin gene transcripts were masked from Cuffdiff, using option -M (see section on “Globin expression estimation” below). Transcript abundance levels pooled across replicates were expressed in terms of log2-transformed FPKMs (Fragments Per Kilobase of exon model per Million mapped fragments), after addition of a value of 1.1 as noise. Noise addition was done to avoid log-transforms of zero values and divide-by-zero issues. Thus, genes with FPKM of 0 are log2-transformed to an expression level of 0.1375. Genes whose transcripts passed the FDR 0.05 threshold and were above our threshold for expression (log2 FPKM > 3) in both cell types being compared were considered to be differentially expressed.

### Globin gene expression estimation

Globin genes are expressed in enormous amounts in erythroid cells. Despite the availability of high-performance compute clusters, estimating abundances for globin mRNAs and performing differential expression tests at these loci is time and memory-intensive, and programs often do not complete running, depending upon the number of reads. To avoid these issues, we (bioinformatically) masked the alpha and beta globin loci on chr11 and chr7, while estimating expression levels and performing differential expression tests for the other genes using Cuffdiff. This is done by supplying a GTF with globin loci to Cuffdiff, using the option -M/--maskfile. As a result, all alpha and beta globin genes, including fetal globins, were reported as not differentially expressed, with an FPKM of 0. To obtain a measure of expression for globins, we extrapolated the expected FPKM of globins from their read counts, by comparing read counts and FPKMs of the top 20 highly expressed genes. The ratio of read counts to FPKMs (∼2 in this case) was used to estimate globin gene expression levels. Therefore, the reported expression level of globin genes is not from Cuffdiff.

### Pseudoreplicates approach for MEG

To avoid the low number of alignments for MEG replicate 2 affecting our analyses and differential expression tests, we used pseudoreplicates for the MEG sample. Pseudoreplicates were generated by pooling alignments from both MEG replicates and randomly splitting the pooled alignments into datasets with equal numbers of alignments. Pseudoreplicate expression levels from Cufflinks (each replicate separately) and Cuffdiff (pooled values from pseudoreplicates) were used for analysis and to perform pairwise differential expression tests. This approach underestimates the real variance in the data, so to avoid potential false discoveries during differential expression testing, we used a more stringent FDR of 0.02 for the identification of differentially expressed genes specifically for the MEP vs. MEG comparison.

### Functional term enrichments

Functional term enrichments were computed using the GREAT tool (46) by using 20 bp upstream of the promoter of each gene as input. Genes from each of the differential expression categories were examined separately, and terms with enrichments at or below a binomial FDR < 0.05 were retained. This produced a list of 1634 terms from multiple ontologies enriched in at least one of the differential expression gene clusters (Supporting Information Table 2). Redundancies in this list were removed to generate a set of 200 terms covering the common themes in the enriched terms. A subset of these terms was extracted to emphasize the five major classes of terms presented in Figure 5A (Supporting Information Table 2).

### Enrichment of occupancy by transcription factors

For each group of genes, we calculated the ratio of % occupancy by a TF in that expression category to the % occupancy by the same TF in the genomic background (all 1239 non-differentially expressed genes). These values were then log2-transformed to represent enrichment (positive values) and depletion (negative values).

### Categorization of cCREs by ATAC-seq signal state across cell types

A set of cCREs was considered associated with a gene, i.e. potentially involved in its regulation, if the elements in that set occurred within a genomic interval extending from 10kb upstream of the transcription start site to 10kb downstream of the polyA addition site of the gene. For all 7,590 genes expressed in at least one of the three cell types, we collected the associated cCREs to generate 48,701 cCRE-gene pairs. In situations of close or overlapping genes, a cCRE could be assigned to more than one gene. We tabulated the mean normalized ATAC-seq signal in mouse blood cell types using information in dbCRE from the VISION project (24). Specifically, we used the negative logarithm (base 10) of the p-value for deviation from a negative binomial distribution as the quantitation for the normalized ATAC-seq signal. The distribution of ATAC-seq signals was evaluated in a subset of the mouse blood cell types, which included the hematopoietic stem and multilineage progenitor cells LSK, CMP, MEP, and GMP, the lineage-restricted progenitor cells CFUMEG and CFUE, and lineage-restricted cells immature MEG and erythroblasts. The distribution of these normalized ATAC-seq signals were compared to the distribution of RNA-seq signals (log2 FPKM) for the polyA+ RNA-seq for MEP, MEG, and ERY in Figure 6D. For all comparisons of patterns of signals between transcriptomes and chromatin accessibility, we used the transcriptome data from polyA+ RNA-seq in mature MEG and chromatin accessibility from ATAC-seq data in immature MEG because no ATAC-seq data were available for mature MEG. The cCRE-gene pairs along with gene expression levels and normalized ATAC-seq signals at the cCREs in each cell type are provided in Supporting information Table 3.

To systematically examine the patterns of chromatin accessibility across cell types, we assigned the ATAC-seq signals in MEP, immature MEG, and ERY to signal states using thresholds derived from the distributions of ATAC-seq signal strengths. We previously set a value of 1.3 (negative logarithm (base 10) of the p-value) as the threshold for calling an ATAC-seq peak (24). Any cCRE with an ATAC-seq signal value below 1.3 in a cell type was assigned to the low (L) signal state in that cell type. The mean of the ATAC-seq signal across all genes and cell types in the current study was 8.6, and this value was chosen to assign cCREs to the mid-range (M, below 8.6) versus high (H, at least 8.6) signal states in each cell type. The 27 possible combinations of H, M, or L signal states in three cell types gave 27 categories of signal state across cell types (Table 1A). Each of the 48,701 cCRE-gene pairs was assigned to an ATAC-seq signal state.

### Tests of predictions from hypotheses to explain discordance in cell type grouping

Predictions of the hypotheses for explaining discordance in cell grouping between RNA and chromatin accessibility measures were tested by evaluating the observed versus expected frequencies of cCRE-gene pairs in informative groups of categories of ATAC-seq signal states (Table 1B). The observed values were the numbers of cCRE-gene pairs in informative signal state categories for the genes in differential expression clusters 10 or 3, and the expected values were the numbers of cCRE-gene pairs in informative signal state categories for the 7,590 genes expressed in any of the three cell types. The data were arranged in 2×2 contingency tables to compare the numbers of cCRE-gene pairs in or not in the informative signal states for genes in a differential expression cluster (observed) versus the numbers of cCRE-gene pairs in or not in the informative signal states for all expressed genes (expected). The odds ratio, 95% confidence interval, and p-values for deviations from expectation were computed using Fisher’s exact test. For testing a prediction from the hypothesized precocious actuation of cCREs in MEP for some ERY-expressed genes, the informative ATAC-seq signal state categories had a higher signal in MEP and ERY than in MEG (Table 1B1). For testing a prediction from the hypothesized loss of accessibility at cCREs in MEP for some MEG-expressed genes, the informative ATAC-seq signal state categories had a low signal in MEP, medium or high in MEG, and any value in ERY (Table 1B2).

### Data availability

Data are available in the Gene Expression Omnibus (GEO) as accessions GSE40522 for RNA-seq, GSE51338 for ChIP-seq data, and GSE143271 and GSE229101 for chromatin accessibility data. These data are also available at the ENCODE Project data portal (https://www.encodeproject.org). Data can be visualized using track hubs on the VISION project website (https://usevision.org) or on a customized browser (http://main.genome-browser.bx.psu.edu).

## Supporting information

This article contains supporting information.

## Funding and additional information

This work was supported by the National Institutes of Health: National Human Genome Research Institute intramural funds (D.M.B.) and grant 1RC2HG005573 (R.C.H.), and National Institute of Diabetes and Digestive and Kidney Diseases grants DK092318 and P30DK090969 (M.J.W.) and DK065806 (R.C.H.); The Roche Foundation for Anemia Research (M.J.W.); the ASH Scholar Award (V.R.P.) and UPenn Measey Fellowship Award (V.R.P.); and the National Science Foundation (OCI-0821527) for instrumentation. The content is solely the responsibility of the authors and does not necessarily represent the official views of the National Institutes of Health. This manuscript is the result of funding in whole or in part by the National Institutes of Health (NIH). It is subject to the NIH Public Access Policy. Through acceptance of this federal funding, NIH has been given a right to make this manuscript publicly available in PubMed Central upon the Official Date of Publication, as defined by NIH.

## References

1. Davidson, E. H., and Erwin, D. H. (2006) Gene regulatory networks and the evolution of animal body plans Science 311, 796–800

2. Kondo, M., Wagers, A. J., Manz, M. G., Prohaska, S. S., Scherer, D. C., Beilhack, G. F. et al. (2003) Biology of hematopoietic stem cells and progenitors: implications for clinical application Annu Rev Immunol 21, 759–806

3. Orkin, S. H., and Zon, L. I. (2008) Hematopoiesis: an evolving paradigm for stem cell biology Cell 132, 631–644

4. Kondo, M., Weissman, I. L., and Akashi, K. (1997) Identification of clonogenic common lymphoid progenitors in mouse bone marrow Cell 91, 661–672

5. Weissman, I. L., Anderson, D. J., and Gage, F. (2001) Stem and progenitor cells: origins, phenotypes, lineage commitments, and transdifferentiations Annu Rev Cell Dev Biol 17, 387–403

6. Iwasaki, H., and Akashi, K. (2007) Myeloid lineage commitment from the hematopoietic stem cell Immunity 26, 726–740

7. Weintraub, H., Campbell Gle, M., and Holtzer, H. (1971) Primitive erythropoiesis in early chick embryogenesis. I. Cell cycle kinetics and the control of cell division J Cell Biol 50, 652–668

8. Hu, J., Liu, J., Xue, F., Halverson, G., Reid, M., Guo, A., et al. (2013) Isolation and functional characterization of human erythroblasts at distinct stages: implications for understanding of normal and disordered erythropoiesis in vivo Blood 121, 3246–3253

9. Hattangadi, S. M., Wong, P., Zhang, L., Flygare, J., and Lodish, H. F. (2011) From stem cell to red cell: regulation of erythropoiesis at multiple levels by multiple proteins, RNAs, and chromatin modifications Blood 118, 6258–6268

10. Gieger, C., Radhakrishnan, A., Cvejic, A., Tang, W., Porcu, E., Pistis, G., et al. (2011) New gene functions in megakaryopoiesis and platelet formation Nature 480, 201–208

11. Deutsch, V. R., and Tomer, A. (2006) Megakaryocyte development and platelet production Br J Haematol 134, 453–466

12. Schulze, H., Korpal, M., Hurov, J., Kim, S. W., Zhang, J., Cantley, L. C. et al. (2006) Characterization of the megakaryocyte demarcation membrane system and its role in thrombopoiesis Blood 107, 3868–3875

13. Debili, N., Coulombel, L., Croisille, L., Katz, A., Guichard, J., Breton-Gorius, J. et al. (1996) Characterization of a bipotent erythro-megakaryocytic progenitor in human bone marrow Blood 88, 1284–1296

14. Klimchenko, O., Mori, M., Distefano, A., Langlois, T., Larbret, F., Lecluse, Y., et al. (2009) A common bipotent progenitor generates the erythroid and megakaryocyte lineages in embryonic stem cell-derived primitive hematopoiesis Blood 114, 1506–1517

15. Pishesha, N., Thiru, P., Shi, J., Eng, J. C., Sankaran, V. G., and Lodish, H. F. (2014) Transcriptional divergence and conservation of human and mouse erythropoiesis Proc Natl Acad Sci U S A 111, 4103–4108

16. An, X., Schulz, V. P., Li, J., Wu, K., Liu, J., Xue, F., et al. (2014) Global transcriptome analyses of human and murine terminal erythroid differentiation Blood 123, 3466–3477

17. Doré, L. C., Chlon, T. M., Brown, C. D., White, K. P., and Crispino, J. D. (2012) Chromatin occupancy analysis reveals genome-wide GATA factor switching during hematopoiesis Blood 119, 3724–3733

18. Doré, L. C., and Crispino, J. D. (2011) Transcription factor networks in erythroid cell and megakaryocyte development Blood 118, 231–239

19. Pimkin, M., Kossenkov, A. V., Mishra, T., Morrissey, C. S., Wu, W., Keller, C. A. et al. (2014) Divergent functions of hematopoietic transcription factors in lineage priming and differentiation during erythro-megakaryopoiesis Genome Res 24, 1932–1944

20. Wu, W., Morrissey, C. S., Keller, C. A., Mishra, T., Pimkin, M., Blobel, G. A. et al. (2014) Dynamic shifts in occupancy by TAL1 are guided by GATA factors and drive large-scale reprogramming of gene expression during hematopoiesis Genome Res 24, 1945–1962

21. Yue, F., Cheng, Y., Breschi, A., Vierstra, J., Wu, W., Ryba, T. et al. (2014) A comparative encyclopedia of DNA elements in the mouse genome Nature 515, 355–364

22. Heuston, E. F., Keller, C. A., Lichtenberg, J., Giardine, B., Anderson, S. M., Center, N. I. H. I. S., et al. (2018) Establishment of regulatory elements during erythro-megakaryopoiesis identifies hematopoietic lineage-commitment points Epigenetics Chromatin 11, 22

23. Xiang, G., Keller, C. A., Heuston, E., Giardine, B. M., An, L., Wixom, A. Q. et al. (2020) An integrative view of the regulatory and transcriptional landscapes in mouse hematopoiesis Genome Res 30, 472–484

24. Xiang, G., He, X., Giardine, B. M., Isaac, K. J., Taylor, D. J., McCoy, R. C. et al. (2024) Interspecies regulatory landscapes and elements revealed by novel joint systematic integration of human and mouse blood cell epigenomes Genome Res 34, 1089–1105

25. Fujiwara, Y., Browne, C. P., Cunniff, K., Goff, S. C., and Orkin, S. H. (1996) Arrested development of embryonic red cell precursors in mouse embryos lacking transcription factor GATA-1 Proc Natl Acad Sci U S A 93, 12355–12358

26. Shivdasani, R. A., Fujiwara, Y., McDevitt, M. A., and Orkin, S. H. (1997) A lineage-selective knockout establishes the critical role of transcription factor GATA-1 in megakaryocyte growth and platelet development Embo J 16, 3965–3973

27. Tijssen, M. R., Cvejic, A., Joshi, A., Hannah, R. L., Ferreira, R., Forrai, A. et al. (2011) Genome-wide analysis of simultaneous GATA1/2, RUNX1, FLI1, and SCL binding in megakaryocytes identifies hematopoietic regulators Dev Cell 20, 597–609

28. Tallack, M. R., Whitington, T., Yuen, W. S., Wainwright, E. N., Keys, J. R., Gardiner, B. B. et al. (2010) A global role for KLF1 in erythropoiesis revealed by ChIP-seq in primary erythroid cells Genome Res 20, 1052–1063

29. Pilon, A. M., Ajay, S. S., Kumar, S. A., Steiner, L. A., Cherukuri, P. F., Wincovitch, S. et al. (2011) Genome-wide ChIP-Seq reveals a dramatic shift in the binding of the transcription factor erythroid Kruppel-like factor during erythrocyte differentiation Blood 118, e139–148

30. Lara-Astiaso, D., Weiner, A., Lorenzo-Vivas, E., Zaretsky, I., Jaitin, D. A., David, E. et al. (2014) Immunogenetics. Chromatin state dynamics during blood formation Science 345, 943–949

31. Corces, M. R., Buenrostro, J. D., Wu, B., Greenside, P. G., Chan, S. M., Koenig, J. L. et al. (2016) Lineage-specific and single-cell chromatin accessibility charts human hematopoiesis and leukemia evolution Nat Genet 48, 1193–1203

32. Nestorowa, S., Hamey, F. K., Pijuan Sala, B., Diamanti, E., Shepherd, M., Laurenti, E., et al. (2016) A single-cell resolution map of mouse hematopoietic stem and progenitor cell differentiation Blood 128, e20-31

33. Laurenti, E., and Göttgens, B. (2018) From haematopoietic stem cells to complex differentiation landscapes Nature 553, 418–426

34. Psaila, B., Barkas, N., Iskander, D., Roy, A., Anderson, S., Ashley, N., et al. (2016) Single-cell profiling of human megakaryocyte-erythroid progenitors identifies distinct megakaryocyte and erythroid differentiation pathways Genome Biol 17, 83

35. Lu, Y. C., Sanada, C., Xavier-Ferrucio, J., Wang, L., Zhang, P. X., Grimes, H. L. et al. (2018) The Molecular Signature of Megakaryocyte-Erythroid Progenitors Reveals a Role for the Cell Cycle in Fate Specification Cell reports 25, 2083–2093 e2084

36. Xavier-Ferrucio, J., and Krause, D. S. (2018) Concise Review: Bipotent Megakaryocytic-Erythroid Progenitors: Concepts and Controversies Stem Cells 36, 1138–1145

37. Pronk, C. J., Rossi, D. J., Mansson, R., Attema, J. L., Norddahl, G. L., Chan, C. K. et al. (2007) Elucidation of the phenotypic, functional, and molecular topography of a myeloerythroid progenitor cell hierarchy Cell Stem Cell 1, 428–442

38. Paralkar, V. R., Mishra, T., Luan, J., Yao, Y., Kossenkov, A. V., Anderson, S. M. et al. (2014) Lineage and species-specific long noncoding RNAs during erythro-megakaryocytic development Blood 123, 1927–1937

39. Parkhomchuk, D., Borodina, T., Amstislavskiy, V., Banaru, M., Hallen, L., Krobitsch, S., et al. (2009) Transcriptome analysis by strand-specific sequencing of complementary DNA Nucleic Acids Res 37, e123

40. Trapnell, C., Williams, B. A., Pertea, G., Mortazavi, A., Kwan, G., van Baren, M. J., et al. (2010) Transcript assembly and quantification by RNA-Seq reveals unannotated transcripts and isoform switching during cell differentiation Nat Biotechnol 28, 511–515

41. Roberts, A., Trapnell, C., Donaghey, J., Rinn, J. L., and Pachter, L. (2011) Improving RNA-Seq expression estimates by correcting for fragment bias Genome Biol 12, R22

42. Trapnell, C., Roberts, A., Goff, L., Pertea, G., Kim, D., Kelley, D. R. et al. (2012) Differential gene and transcript expression analysis of RNA-seq experiments with TopHat and Cufflinks Nat Protoc 7, 562–578

43. Trapnell, C., Hendrickson, D. G., Sauvageau, M., Goff, L., Rinn, J. L., and Pachter, L. (2013) Differential analysis of gene regulation at transcript resolution with RNA-seq Nat Biotechnol 31, 46–53

44. Eichler, G. S., Huang, S., and Ingber, D. E. (2003) Gene Expression Dynamics Inspector (GEDI): for integrative analysis of expression profiles Bioinformatics 19, 2321–2322

45. Mortazavi, A., Pepke, S., Jansen, C., Marinov, G. K., Ernst, J., Kellis, M. et al. (2013) Integrating and mining the chromatin landscape of cell-type specificity using self-organizing maps Genome Res 23, 2136–2148

46. McLean, C. Y., Bristor, D., Hiller, M., Clarke, S. L., Schaar, B. T., Lowe, C. B. et al. (2010) GREAT improves functional interpretation of cis-regulatory regions Nat Biotechnol 28, 495–501

47. Wilson, N. K., Foster, S. D., Wang, X., Knezevic, K., Schütte, J., Kaimakis, P. et al. (2010) Combinatorial transcriptional control in blood stem/progenitor cells: genome-wide analysis of ten major transcriptional regulators Cell Stem Cell 7, 532–544

48. Soler, E., Andrieu-Soler, C., de Boer, E., Bryne, J. C., Thongjuea, S., Stadhouders, R. et al. (2010) The genome-wide dynamics of the binding of Ldb1 complexes during erythroid differentiation Genes Dev 24, 277–289

49. Shearstone, J. R., Pop, R., Bock, C., Boyle, P., Meissner, A., and Socolovsky, M. (2011) Global DNA demethylation during mouse erythropoiesis in vivo Science 334, 799–802

50. Akashi, K., He, X., Chen, J., Iwasaki, H., Niu, C., Steenhard, B. et al. (2003) Transcriptional accessibility for genes of multiple tissues and hematopoietic lineages is hierarchically controlled during early hematopoiesis Blood 101, 383–389

51. Mansson, R., Hultquist, A., Luc, S., Yang, L., Anderson, K., Kharazi, S. et al. (2007) Molecular evidence for hierarchical transcriptional lineage priming in fetal and adult stem cells and multipotent progenitors Immunity 26, 407–419

52. Efroni, S., Duttagupta, R., Cheng, J., Dehghani, H., Hoeppner, D. J., Dash, C. et al. (2008) Global transcription in pluripotent embryonic stem cells Cell Stem Cell 2, 437–447

53. Huang, H., and Cantor, A. B. (2009) Common features of megakaryocytes and hematopoietic stem cells: what’s the connection? J Cell Biochem 107, 857–864

54. Trapnell, C., Pachter, L., and Salzberg, S. L. (2009) TopHat: discovering splice junctions with RNA-Seq Bioinformatics 25, 1105–1111

55. Kim, D., Pertea, G., Trapnell, C., Pimentel, H., Kelley, R., and Salzberg, S. L. (2013) TopHat2: accurate alignment of transcriptomes in the presence of insertions, deletions and gene fusions Genome Biol 14, R36

56. Cabili, M. N., Trapnell, C., Goff, L., Koziol, M., Tazon-Vega, B., Regev, A. et al. (2011) Integrative annotation of human large intergenic noncoding RNAs reveals global properties and specific subclasses Genes Dev 25, 1915–1927

57. Li, H., Handsaker, B., Wysoker, A., Fennell, T., Ruan, J., Homer, N. et al. (2009) The Sequence Alignment/Map format and SAMtools Bioinformatics 25, 2078–2079

58. Quinlan, A. R., and Hall, I. M. (2010) BEDTools: a flexible suite of utilities for comparing genomic features Bioinformatics 26, 841–842

59. Karolchik, D., Hinrichs, A. S., Furey, T. S., Roskin, K. M., Sugnet, C. W., Haussler, D., et al. (2004) The UCSC Table Browser data retrieval tool Nucleic Acids Res 32, D493–D496

60. Kent, W. J., Sugnet, C. W., Furey, T. S., Roskin, K. M., Pringle, T. H., Zahler, A. M. et al. (2002) The human genome browser at UCSC Genome Res 12, 996–1006.

61. Kuhn, R. M., Haussler, D., and Kent, W. J. (2013) The UCSC genome browser and associated tools Brief Bioinform 14, 144–161

62. Kent, W. J., Zweig, A. S., Barber, G., Hinrichs, A. S., and Karolchik, D. (2010) BigWig and BigBed: enabling browsing of large distributed datasets Bioinformatics 26, 2204–2207

63. Meyer, L. R., Zweig, A. S., Hinrichs, A. S., Karolchik, D., Kuhn, R. M., Wong, M., et al. (2013) The UCSC Genome Browser database: extensions and updates 2013 Nucleic Acids Res 41, D64–69

