## Supporting Information: Supplemental Figures for "Divergence between transcriptomes and chromatin accessibility during differentiation from a bipotential progenitor cell population to erythroblasts and megakaryocytes"

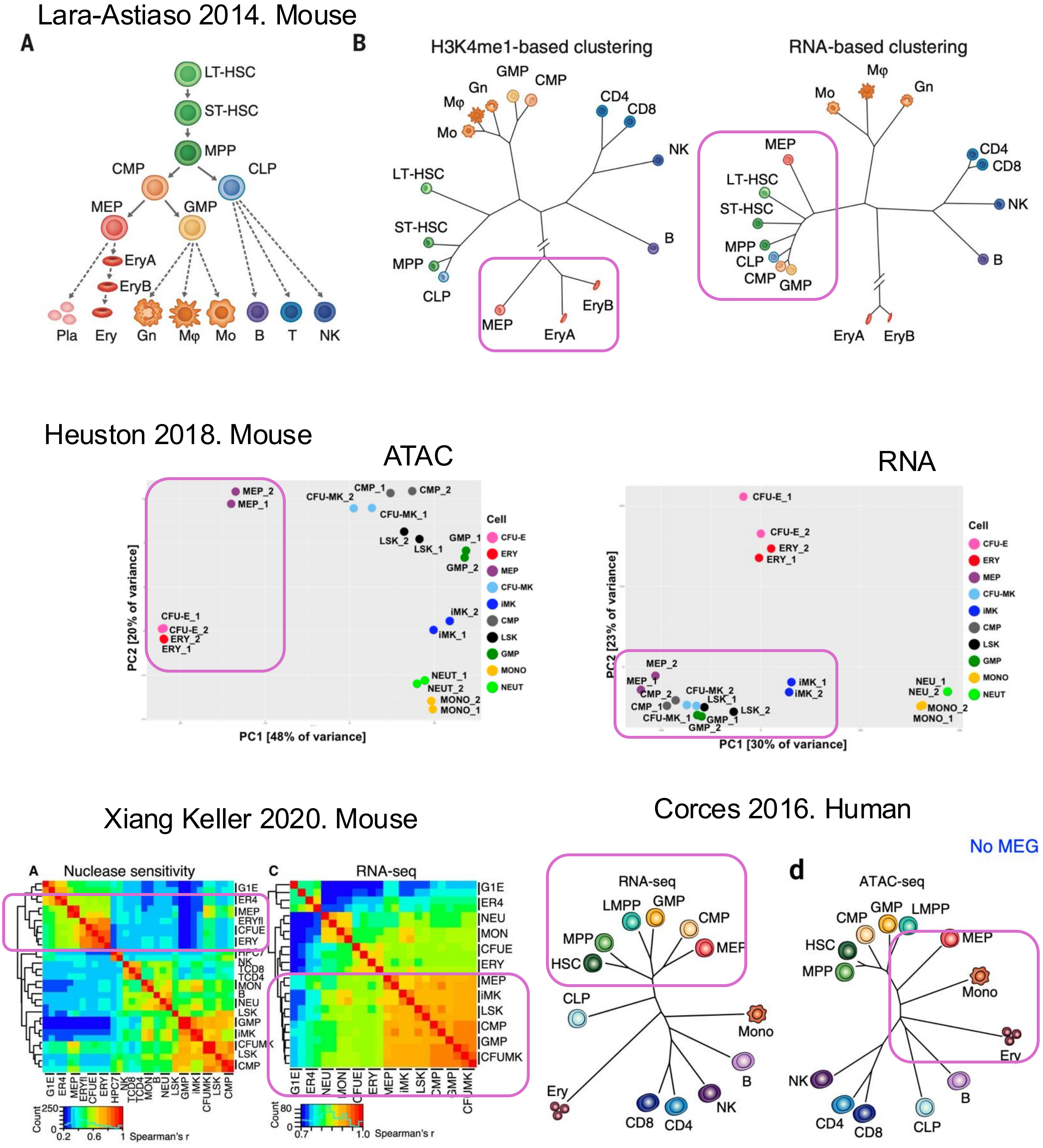


**Supplemental Figure S1.** Published results showing MEP grouping with MEG and multilineage progenitor cells when using RNA levels as a distance metric but MEP grouping with ERY when using chromatin accessibility or histone modification (H3K4me1) as the distance metric. These images were taken from the published figures and annotated with rounded rectangles to emphasize the discordance. These results were synthesized in the summary shown in Figure 1A. The papers are Lara-Astiaso 2014 (reference 30), Heuston 2018 (reference 22), Xiang Keller 2020 (reference 23), and Corces 2016 (reference 31).

A

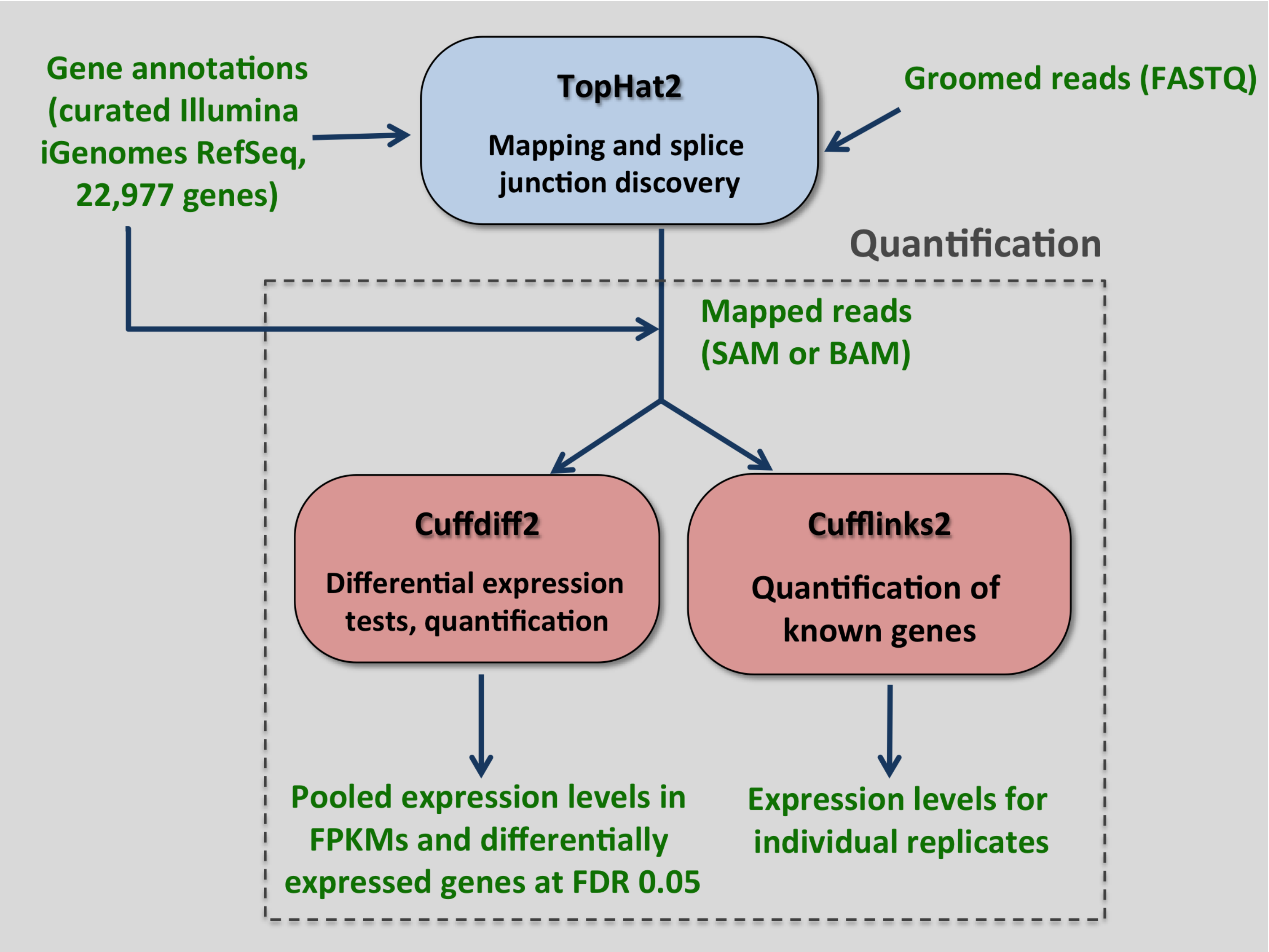


B

| | # Read Pairs | # Mapped Reads | # Alignments | % Mapped Reads |
|---|---|---|---|---|
| MEP Rep1 | 2 x 123,922,076 | 94,308,729 | 180,399,326 | 76.1% |
| MEP Rep2 | 2 x 120,129,876 | 84,178,334 | 161,120,323 | 70.1% |
| ERY Rep1 | 2 x 117,566,409 | 61,078,428 | 115,103,757 | 52% |
| ERY Rep2 | 2 x 106,890,312 | 49,524,755 | 89,069,700 | 46.3% |
| MEG Rep1 | 2 x 107,938,301 | 58,026,025 | 112,235,879 | 53.8% |
| MEG Rep2 | 2 x 101,667,130 | 3,491,255 | 7,332,882 | 3.4% |

**Supplemental Figure S2. RNA-seq data processing** (A) Schematic of the RNA-seq analysis pipeline. Tools used are in colored, solid boxes. Input and output files and data are indicated in green. (B) RNA-seq read mapping statistics. For each sample, the number of sequenced reads, mapped reads, and alignments, as well as percentage of mapped reads are shown.
Col. 1: # Read Pairs = number of paired sequence reads, where each read in a pair has the same read ID
Col. 2: # Mapped Reads = number of unique read IDs with at least one end mapped
Col. 3: # Alignments = number of genomic locations ("hits to the genome") the reads mapped to, when allowing multiple mapping.
Col. 4: % Mapped Reads = (Col. 2/Col.1)*100

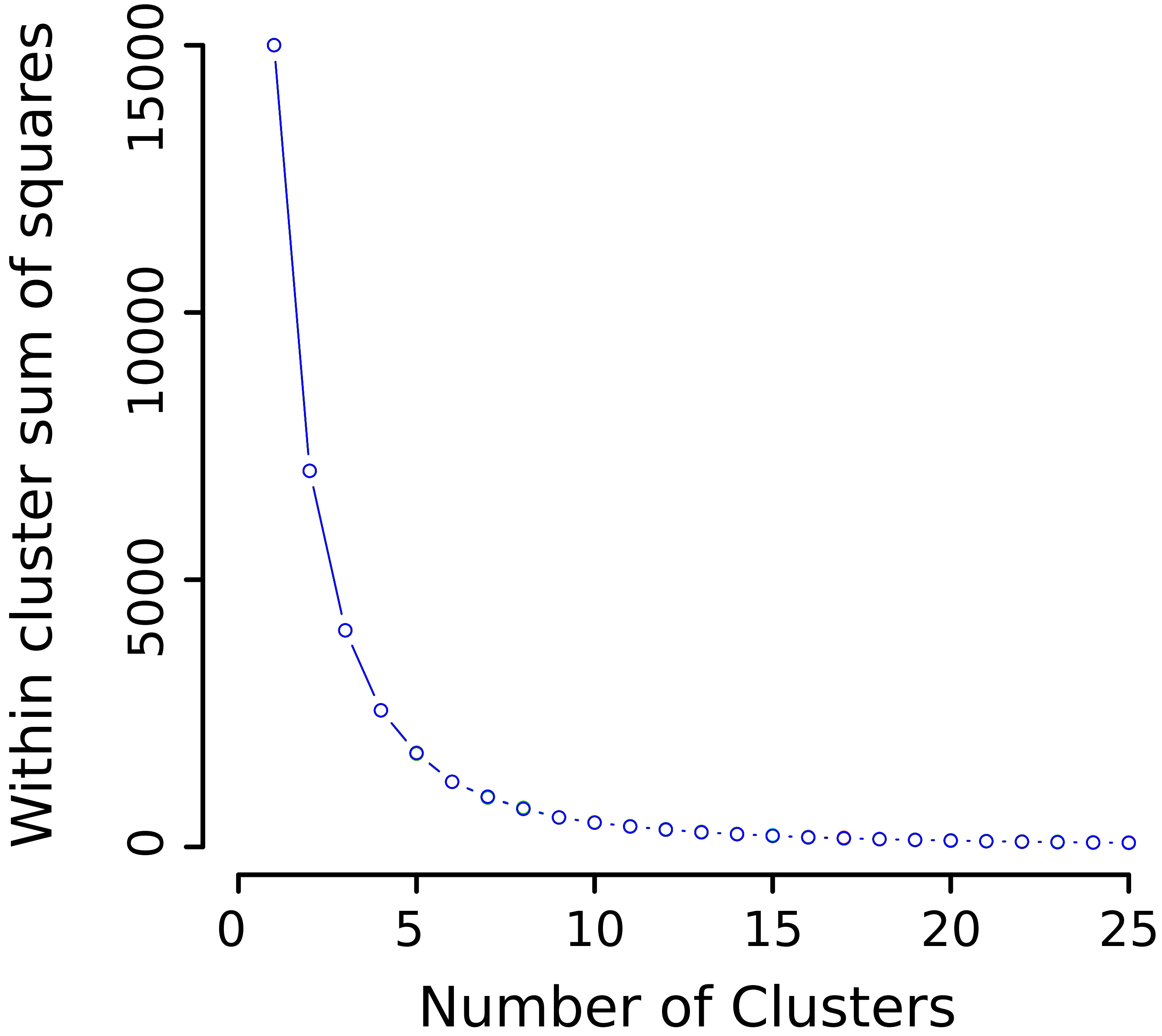


**Supplemental Figure S3. Choice of "k" or number of clusters.** K-means clustering was performed on standardized gene e pression levels for k ranging from 1 to 25, and the within cluster sum-of-squares (WCSS) was calculated for each value of "k". This procedure was performed 20 times. The plot shows the WCSS (hollow circles) obtained at each value of "k", for each of the 20 iterations. The "k" at which the within cluster sum-of-squares did not decrease drastically with an increase in number of clusters (k = 12) was chosen as the optimal number of groups to partition genes into.

| TF | # Peaks | Cell type | Source |
|---|---|---|---|
| GATA1 | 5767 | TER119+ fetal liver erythroblasts | Pimkin et. al. |
| TAL1 | 3086 | TER119+ fetal liver erythroblasts | Pimkin et. al. |
| ETO2 | 1455 | DMSO Induced MEL | Soler et. al. |
| MTGR1 | 4682 | DMSO Induced MEL | Soler et. al. |
| GATA1 | 1727 | CD41+ fetal liver megakaryocytes | Pimkin et. al. |
| GATA2 | 2728 | CD41+ fetal liver megakaryocytes | Pimkin et. al. |
| TAL1 | 3505 | CD41+ fetal liver megakaryocytes | Pimkin et. al. |
| FLI1 | 2001 | CD41+ fetal liver megakaryocytes | Pimkin et. al. |
| GATA2 | 9099 | HPC-7 | Wilson et. al. |
| LMO2 | 9006 | HPC-7 | Wilson et. al. |
| LYL1 | 4196 | HPC-7 | Wilson et. al. |
| RUNX1 | 4988 | HPC-7 | Wilson et. al. |
| TF Heptad | 1007 | HPC-7 | Wilson et. al. |

**Supplemental Figure S4: Peak statistics and sources of TF occupancy data**
